# Evolutionary dynamics of nascent multicellular lineages

**DOI:** 10.1101/2024.05.10.593459

**Authors:** Guilhem Doulcier, Philippe Remigi, Daniel Rexin, Paul B. Rainey

## Abstract

The evolution of multicellular organisms involves the emergence of cellular collectives that eventually become units of selection in their own right. The process can be facilitated by ecological conditions that impose heritable variance in fitness on nascent collectives with long-term persistence depending on capacity of competing lineages to transition reliably between soma- and germ-like stages of proto-life cycles. Prior work with experimental bacterial populations showed rapid increases in collective-level fitness with capacity to switch between life cycle phases being a particular focus of selection. Here we report experiments in which the most successful lineage from the earlier study was further propagated for 10 life cycle generations under regimes that required different investment in the soma-like phase. To explore the adaptive significance of switching, a control was included in which reliable transitioning between life cycle phases was abolished. The switch proved central to maintenance of fitness. Moreover, in a non-switch treatment, where solutions to producing a robust and enduring soma-phase were required, evolution of *mutL*-dependent switching emerged *de novo*. A newly developed computational pipeline (`colgen`) was used to display the moment-by-moment evolutionary dynamics of lineages providing rare visual evidence of the roles of chance, history and selection. `Colgen`, underpinned by a Bayesian model, was further used to propagate hundreds of mutations back through temporal genealogical series, predict lineages and time points corresponding to changes of likely adaptive significance, and in one instance, via a combination of targeted sequencing, genetics and analyses of fitness consequences, adaptive significance of a single mutation was demonstrated. Overall, our results shed light on the mechanisms by which collectives adapt to new selective challenges and demonstrate the value of genealogy-centered approaches for investigating the dynamics of lineage-level selection.

## Introduction

Major evolutionary transitions in individuality underpin the emergence of life’s complexity (1–4). A transition of particular interest is that involving the formation of multicellular collectives from ancestral unicellular types (5–8). The transition initiates with the formation of simple groups – for example, through failure of cells to separate upon cell division – and completes once groups become units of selection in their own right. For the latter, newly formed collectives must be Darwinian, that is, collectives must be discrete (with variation among collectives), replicate, and leave offspring copies that resemble parental types (9).

As elaborated elsewhere, these Darwinian properties are derived traits and require evolutionary explanation (10, 11). But how these properties evolve – in the absence of collectives already manifesting these properties – has been a puzzle. One solution that has motivated experimental studies recognises that variation, replication and heredity can be exogenously imposed (scaffolded) by specific ecological conditions (12–16). Once stably imposed, simple undifferentiated groups of cells can become unwitting players in a selective process that occurs over timescales defined by the birth and death of groups, just as if they were cells of a multicellular organism.

A conceivable ecological setting – and one that has inspired laboratory studies – is provided by reeds in a pond (12). Reeds act as scaffolds around which microbial mats attach allowing colonisation of the air-liquid interface (ALI), but importantly spatial separation of reeds ensures that mats are discrete, thus allowing variation to manifest at the level of mats. Periodically mats fail and sink, opening niches for formation of new mats. If dispersing cells from an extant mat re-colonise a vacant reed, then the newly formed mat, being derived from the extant parent, may inherit parental characteristics. Mats thus become units of selection in their own right with selection operating over two timescales: the doubling time of mats, and the doubling time of cells (13). Knowledge of the evolutionary dynamics that unfold at the level of mats stands to shed light on processes underpinning the origin of multicellular life.

In the laboratory, an analogy of the reeds-in-pond model has been devised by substituting glass vials for reeds with experimental realisation in earlier studies (17, 18). In brief, the work uses the aerobic bacterium *Pseudomonas fluorescens* SBW25, whose metabolic activities, when introduced into static broth microcosms, causes the medium to become anoxic (19). This creates conditions that favour cellulose-over-producing, ALI-colonising (mat-forming) mutants known as “wrinkly spreaders” (WS). Presence at the ALI ensures that constituent cells benefit from freely available oxygen. Given a cost associated with the over-production of cellulose, WS mats are susceptible to invasion by cellulose defective “smooth” (SM) mutants. These types pay none of the costs, but fail to provide structural support to the mat, ultimately lead to mat collapse (20). While this amounts to a classic tragedy of the commons (21), taken back into the pond setting, functional roles of WS mat-forming cells and swimming SM cells can be viewed as analogous to primitive soma and germ phases, respectively (11). WS mats perform an ecological role (ensuring access to oxygen) while producing seeds (SM cells) of the next generation of mats. The latter is especially relevant because in the absence of propagules, mats, like soma, are an evolutionary dead-end (17).

A graphical representation of both the life cycle and the lineage selection regime is provided in Figure 1. Negative frequency dependent interaction among WS and SM cells generates a life cycle that in appropriate ecological context becomes the primary focus of selection. Important to note are the causes of lineage-level death-birth dynamics. Lineages go extinct whenever they fail to generate the next stage in the life cycle, or because the mat phase collapses prematurely (soma failure). Given that the life cycle is fuelled by spontaneous mutation, lineages are particularly prone to failure, with lineage death providing opportunity for extant lineages (chosen at random) to export their reproductive success to new microcosms in precisely the same way as a mat that falls from a reed provides opportunity for competing mats to reproduce.

**Figure 1.**
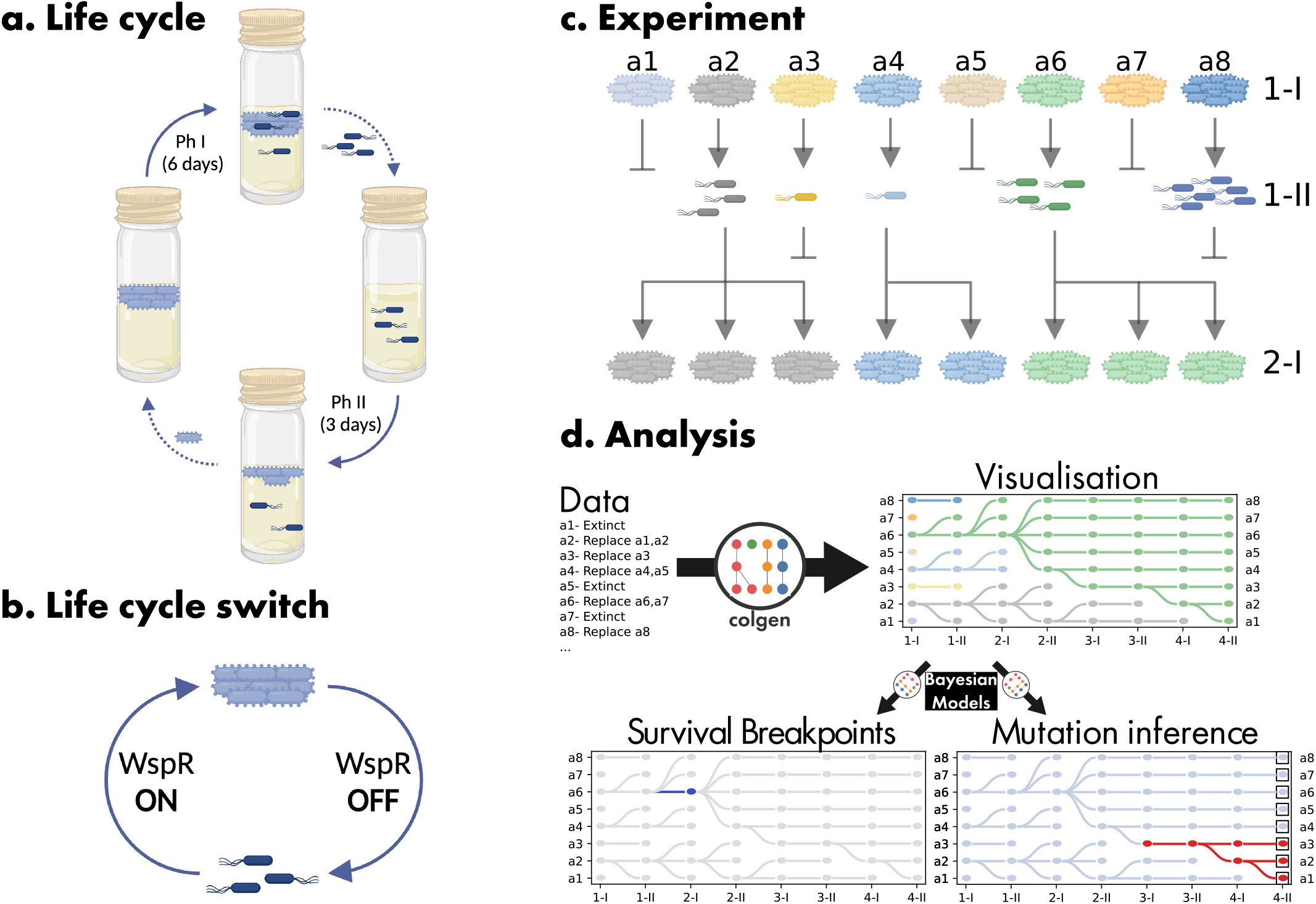
Experimental regime and colgen-based representation. **a** A single generation of the nascent multicellular life cycle has two phases. Phase I (Ph I) initiates with placement of a single WS genotype into a glass microcosm. The WS type over-produces cellulose and forms a soma-like structure (a mat) at the air-liquid interface (ALI). During the six day Ph I stage the mat, in addition to remaining at the ALI, must generate non-cellulose producing (SM) cells that function as seeds (germ cells) for the next generation of mats (solid arrow). At the end of Ph I, SM germ cells are transferred to a fresh microcosm (dotted arrow) where the soma state must be re-established. After three days in Phase II (Ph II) WS cells are collected and a single genotype used to found the next generation. Alternative experimental setups are described and contrasted in (17, 18). **b** Transition between soma (WS) and germ (SM) states is required for survival and involves activation and deactivation of cellulose production, which is effected by mutations that activate and deactivate the synthesis of cyclic-di-GMP. Of particular relevance is the derived “life cycle switch” LCS^+^ genotype that has enhanced capacity to transition between soma and germ states because of a mutation in *mutS* that causes frameshifting to occur at high frequency in a tract of guanine (G) residues in the active site of the WspR-encoded diguanylate cyclase. When the reading frame is intact, the cellulose-overproducing soma state is acheived, but gain or loss a G-residue causes a frameshift, deactivation of WspR leading to realisation of the germ state. **c** Extinction of lineages occurs if there are no soma cells after six days of Ph I, no germ cells after three days in Ph II, or if the mat collapses during Ph I (soma failure). Death of lineages allows a death-birth dynamic to unfold with lineages functioning as units of selection in their own right. A single meta-population of eight distinct lineages is shown. While lineages compete, then never mix. Extant lineages replace themselves, but also have the possibility of reproducing on death of a competing lineage. Selection rewards persistence which depends on reliable transitioning between states. **d** Schematic description of three functions of the `colgen` package: visualisation (Section A), survival breakpoint detection (Section B) and belief propagation (Section C).

The experiment originally conducted by (17) was performed over 10 life cycle generations with selection rewarding lineages that repeatedly transitioned through soma and germ phases. On conclusion of the experiment a single lineage (Line 17) had swept to fixation. Detailed analysis showed that success was a consequence of collective-level benefits arising from a simple genetic switch. Of a numerous array of mutations, two were of special significance: one that caused the WspR digualnylate cyclase (DGC) to achieve a state of constitutive activation (thus over-producing c-di-GMP and eliciting over-production of cellulose), and a second, loss-of-function mutation, in *mutS* (A1489C), a gene involved in methyl-directed mismatch repair. The latter, while causing an overall elevation in mutation rate, specifically increases frame-shifting in tracts of guanine (G) residues (22). While tracts of Gs are rarely found in any genome, one – composed of seven G residues – happens to reside in the active site of the DGC domain of WspR. Given the defect in *mutS*, WspR gained ability to switch rapidly between functional and non functional states (due to elevated rates of frame-shifting), thus ensuring reliable transitions between soma and germ phases (17).

Here we took the adaptive genotype and an isogenic lineage in which the genetic switch was rendered inoperable and asked whether the genetic switch delivers longer-term evolutionary advantage. We also explored the extent to which the switch facilitates lineage-level responses to the challenge of growing in larger diameter microcosms requiring substantially greater investment in the soma phase. Underpinning our analyses is `colgen`, a newly developed analytical tool that provides graphical representation of genealogies, a variety of means for exploring lineage dynamics, and, supported by a Bayesian model, allows inferences as to branch points of likely adaptive significance. Further, when combined with DNA sequence from derived lineages `colgen` predicts the sequence of causal mutational events. Subsequent genetic analyses support the value of these predictions.

## Material and Methods

### Experimental evolution of collectives

The experimental protocol is as described previously (17) and depicted in Figure 1a-c. The two genotypes used here, Line 17 (hereafter Life Cycle Switch +, LCS^+^) and the isogenic variant in which the *mutS* A1489C was reverted to wild type (hereafter Life Cycle Switch -, LCS^-^), are derived from (17).

Both LCS^+^ and LCS^-^ genotypes were propagated through 10 life cycle generations in standard (S) 17 mm diameter microcosms, and in parallel, in large (L) microcosms of 35 mm diameter, with respectively 9.5 and 31.5 ml media to keep surface area to volume ratio constant. Two genotypes, multiplied by two sizes of microcosm, results in four experimental regimes, designated S-LCS^+^, L-LCS^+^, S-LCS^-^, L-LCS^-^.

Ten life cycle generations were performed on 48 parallel cultures blocked as six meta-populations of eight lineages. As in the reeds-in-pond thought experiment, lineages that fail to complete the life cycle and go extinct provide opportunity for viable collective lineages to reproduce. Upon demise of a lineage, a viable type from the same block is chosen at random and allowed to replace the extinct type. In the case where all cultures from a given block go extinct at the same time, random cultures from viable blocks are used to replace the extinct lineages. The eight-microcosm sub-population structure is a practical consequence of the experimental setup in eight-microcosm racks. It limits global competition between lineages as subpopulations must be entirely extinct before being invaded by lineages from other meta-populations

### Measuring lineage-level fitness

The standard assay of fitness in experimental evolution is to determine the growth rate of derived genotypes in direct competition with ancestral types ((23), performed for this system in (17)). This captures two components of fitness: fecundity and viability. Here, however, this measure is inadequate. Firstly, it is necessary to consider a timescale longer than one growth period within one microcosm (15) – survival probability over the timescale of collective-generations is relevant for assessing collective level adaptation. Secondly, standard measures capture two components of fitness: fecundity and viability. Observation of the genealogies of lineages as shown in Figure 3 led to realisation that the probability of avoiding extinction is a sound predictor of future evolutionary success (24). In addition, viability (and not fertility) is the only component of fitness that depends on the genotype of collectives in this experimental setup. Indeed, surviving collectives have equal chance of reproduction (with the total number of offspring being set by the number of extinct competing lineages). Henceforth, measures of fitness were determined, not by competition with an ancestral type, but by measuring the number of lineage extinction events after replicate lineages of single test genotypes have gone through a further life cycle generation. Further details are available in Supplementary note 1 including additional growth experiments that show a response to collective-level selection.

### Sequencing

All WS genotypes chosen at the end of generation 10 – in lieu of founding generation 11 – were subject to whole genome sequencing. Bacterial genomic DNA was extracted from static overnight cultures of WS clones using the Wizard Genomic DNA purification kit (**A1125**, Promega) following the manufacturer’s instructions. Whole genome sequencing was performed using Illumina sequencing technology. Short reads were mapped on the *Pseudomonas fluorescens* SBW25 reference genome (GCF000009225_2) and mutations were identified with Breseq v. 0.32.1 (25) using default parameters. When taking into account extinctions within cycle 10 and filtering out sequence data of poor quality, 45 of 48 endpoint genotypes were sequenced for L-LCS^+^, 48 of 48 for S-LCS^+^ 42 of 48 for L-LCS^-^ and 37 of 48 for S-LCS^-^. Further genotypes were targeted for sequencing using the same protocol after the experiment concluded, based on analysis of genealogies. In S-LCS^+^, genotypes at each cycle, in L-LCS^+^ 1-II-A7 (cycle 1, Ph II, lineage A7) 2-I-A5, 2-II-A6, 2-II-A7, 2-II-A8, 3-II-A1 and 3-II-E2 were sequenced in order to elucidate the origin of the *mutL* mutation. Further details are available in Supplementary note 3.

### Analysis of genealogies using the colgen package

Each new cycle of the experiment produced a set of complex data: for each lineage, the record of germ and soma-like cell census, the presence of the soma state, and additionally death-birth events, were recorded. Plotting and visualisation features of `colgen` (26, v 2.0b1) were used to synthesise data without discounting complexity. Survival probability, as determined via `colgen` (Section B, described in Supplementary note 2) was combined with DNA-sequence information to identify genotypes of likely adaptive significance (Section C (described in Supplementary note 3).

All results obtained using `colgen` in the context of this manuscript can be explored interactively online: 10.5281/zenodo.11170876.

## Results

### A. Eco-evolutionary dynamics

Figure 2 shows the frequency of lineage extinctions and their causes. Extant lineages are in blue, with extinctions due to failure to produce germ cells shown in red, failure of the soma phase (mat) in orange, and failure to produce soma cells in green (see figure caption). The proportion of soma failure (i.e., mat collapse) is higher in large microcosms (L) than in standard microcosms (40/96, vs 3/96 in the first cycle). This shows that large microcosms present a new and more stringent challenge to lineages when compared to the ancestral environment of standard microcosms. LCS^+^ treatments showed little change (S-LCS^+^ even less than L-LCS^+^),which indicates that these lineages are close to an evolutionary equilibrium. In contrast, the proportion of lineages that survived in the LCS^-^ treatments was initially low, but rapidly increased. Figure S1 shows the proportion of lineages that avoided extinction at each of the 10 life cycle generations. It also reveals how changes in survival during one phase correlates with changes in survival during the other. Viewed as displayed in Figure S1, and comparing the first and the tenth generations, the data indicate, for S-LCS^-^ and L-LCS^+^, a trade-off between survival in Ph I versus Ph II. For S-LCS^+^, there is a decrease in phase I survival that is not accompanied by an increase in phase II survival. However, phase II survival was already close to its maximal possible value, indicating the founding genotype was close to optimality, thus obscuring any tradeoff. Finally, for L-LCS^-^ no tradeoff is seen, suggesting the series of mutations may have broken the tradeoff (15).

**Figure 2.**
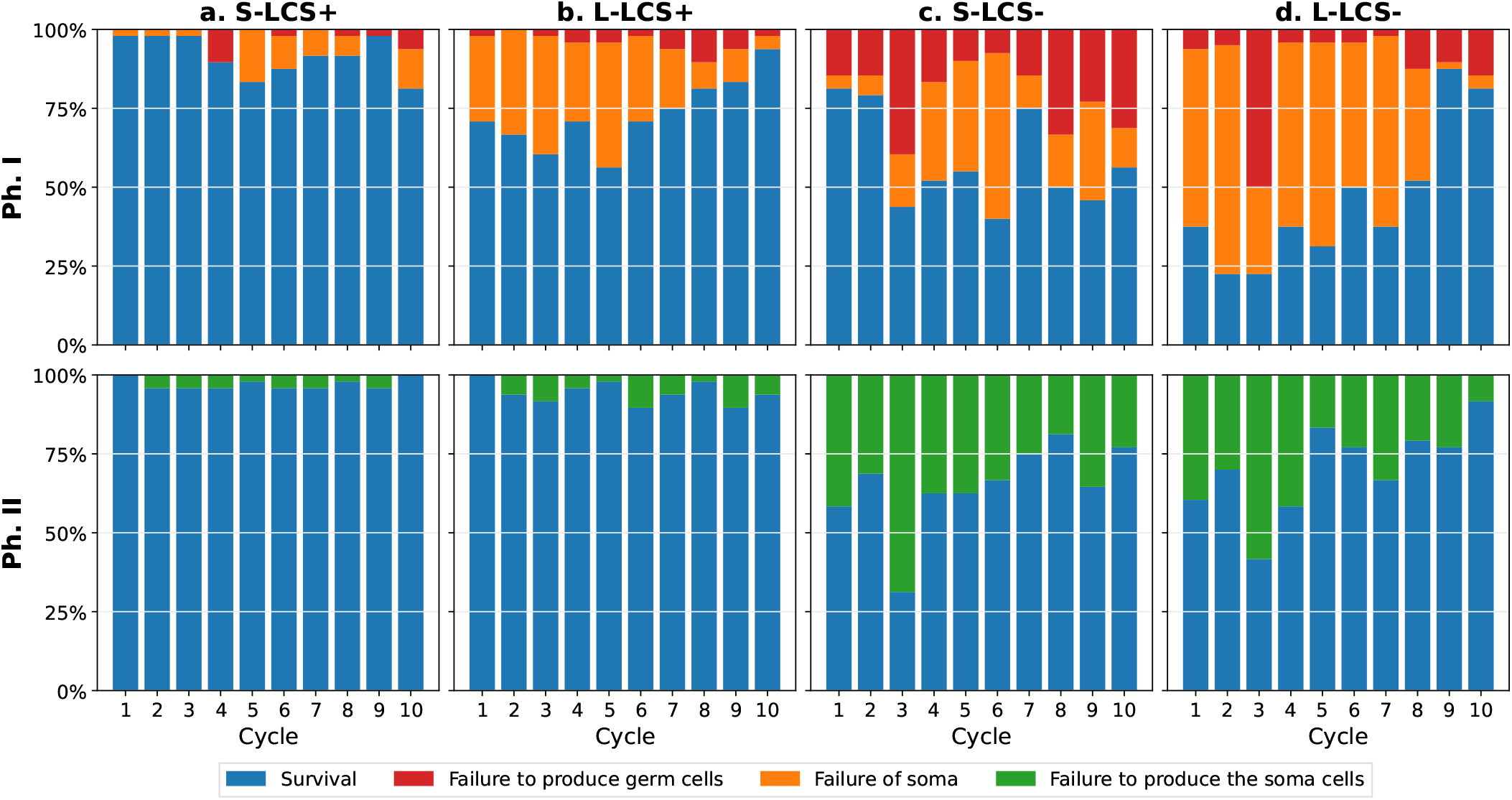
Proportion of surviving collectives in phase I and II for each cycle. The stacked bar correspond to the proportion of surviving collective for each cycle. Blue denotes extant lineages, with red, orange and green marking extinct lineages and causes of extinction: during Ph I, failure to produce the germ cells (red), failure of the mat to remain intact for the required six days (orange), and during Ph II, failure to produce soma cells (green). Each column is different treatment, LCS^+^: Lineage 17 from (17) with a life cycle switch, LCS^-^: same lineage with the switch reverted, S: standard microcosms, L: large microcosms.

Data presented in Figure 2 averages the proportion of collectives that avoid extinction across the entire population of lineages at each time point. It masks heterogeneity, including the fact that some lineages are evolutionary dead-ends, while others rise to fixation. A more information-rich representation is one where the evolutionary fate of each lineage is displayed along with the causes of extinction.

Figure 3 provides such a visual representation. Immediately apparent are treatment-specific differences in lineage dynamics, with those founded by LCS^+^ suffering fewer extinction events compared to those founded by LCS^-^. This is true for both continued selection in standard size microcosms and in the face of challenges posed by evolving in large diameter microcosms. Where improvement is evident, it is most apparent in the capacity of lineages to form mats able to endure the entire six-day period of Ph I. Interestingly this trend is reversed in S-LCS^+^ where death due to mat failure increased during course of the experiment. Production of the germ-line phase during Ph I and soma-phase during Ph II posed greatest challenge to LCS^-^ treatments, but also showed improvement during the course of the experiment.

**Figure 3.**
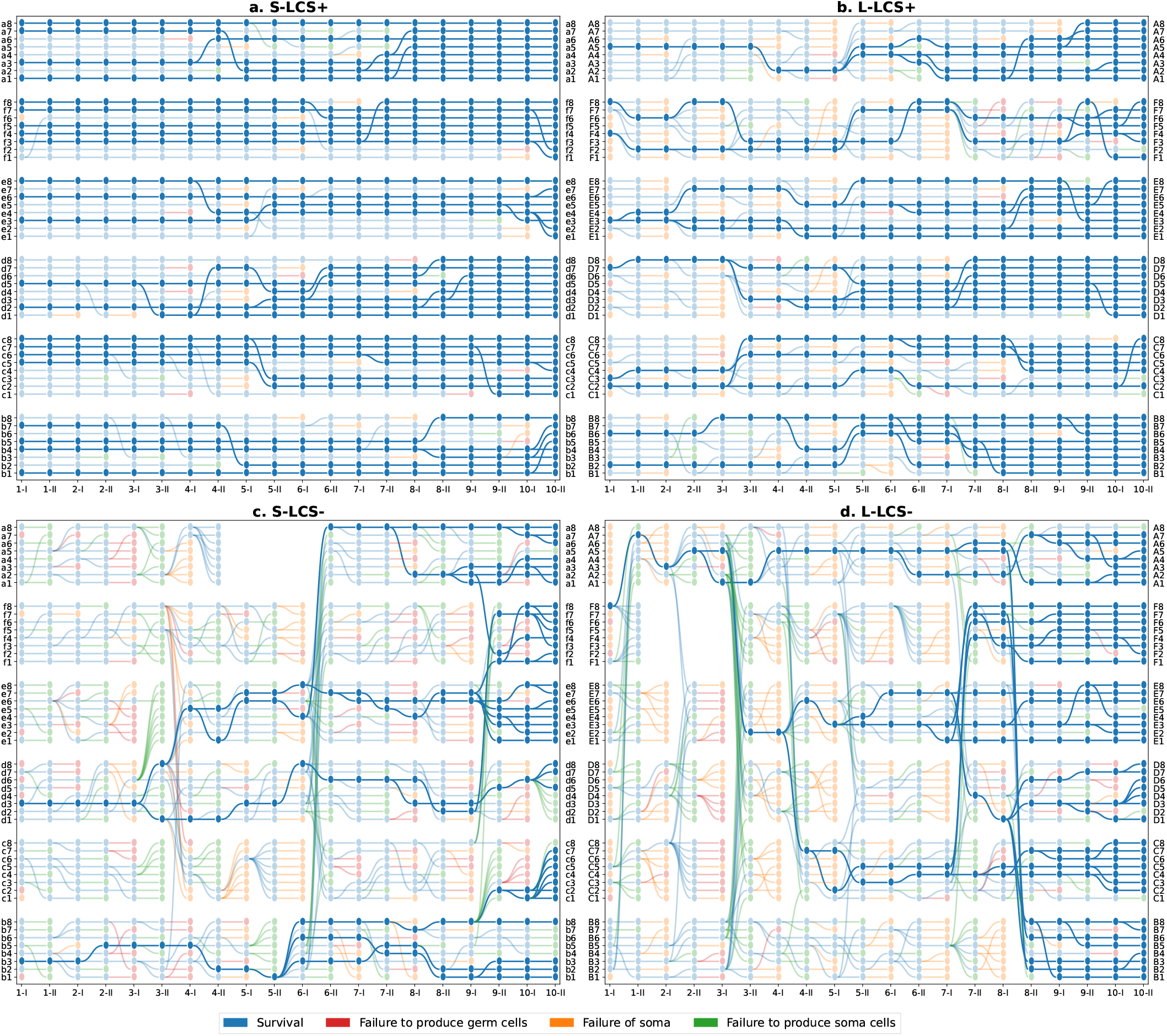
Lineage genealogies. The lineage-level selection experiment yields a rich data set of 48 parallel microcosms (lineages) for each treatment with associated extinction (colours) and replacement events (edges). Lineage-level genealogies in the shape of directed acyclic graphs offer a synthetic but more granular visualisation compared to the proportion of events at each phase (Figure 2). The x-axis labels are phases within each life cycle generation, with 1-I indicating the end of Ph I of the first cycle, 1-II, the end of Ph II of the first cycle and so forth. The gaps (L-LCS^+^ population F in 2-I, 2-II, 3-I and S-LCS^-^ population a in 5-I, 5-II, 6-I) are missing due to technical errors.

Examination of the coalescent tree(s) (depicted in solid colour) reveals further differences among treatments. LCS^+^ maintained in standard microcosms showed few deaths with half of the founding lineages remaining intact at the end of the experiment. On the other hand, LCS^-^ lineages propagated in large microcosms narrowly escaped extinction, with extant lineages, on conclusion of the experiment, being descendants of just a single ancestral microcosm (microcosm 2-II-A5 — generation 2, Ph II, microcosm A5).

Collective fitness was assessed by estimating the probability of a collective to avoid extinction (See Material and Methods). The data, shown in Figures S2 to S5, is largely concordant with that shown in Figure 2. LCS^+^ lineages evolving in standard microcosms showed an overall decline in fitness relative to the ancestral type (Figure S2a) with negative contributions coming from both reductions in durability of the soma phase (Figure S2b) and in capacity to switch between life cycle phases (Figure S2c-d). This negative trend was not apparent for LCS^+^ types in large microcosms, where the fitness of all derived lineages exceed the ancestral type (Figure S3a).

The fitness of derived LCS^-^ lineages in standard microcosms was highly variable, with some test genotypes having elevated fitness, but others with reduced fitness. Low probability of switching was the primary cause of low fitness (Figure S4cd). This is understandable in terms of the availability of mutational routes for switching between soma and germ phases becoming increasingly limited as a consequence of prior mutational cycles. In contrast, in large microcosms, there was a noticeable improvement in fitness of all tested LCS^-^ lineages (Figure S5a), primarily in terms of ability to produce an enduring soma phase (Figure S5b).

### B. Identifying likely adaptive events

The genealogical connections depicted in Figure 3 show what appear to be selective sweeps. Those microcosms that mark the start of such sweeps indicate points at which adaptive mutations may have arisen. However, as the number of offspring at any time-point is dependent on death of competing lineages, the connection to adaptive potential can be difficult to infer. For example, it is not inconceivable that a poorly performing, but nonetheless extant lineage, happens to amplify, not because of improvements in fitness, but by chance. To distinguish improvements in future representation due to chance versus selection, a measure of enduring “adaptive potential” associated with each node is desirable.

To this end, we employed a Bayesian network model in conjunction with an evidence-propagation algorithm (27) to predict improvement in the probability of survival. The model assumes existence of a hidden *probability of survival* value for each node, which is calculated by computing the maximum-a-posteriori value, using an expectation-maximisation algorithm that takes genealogical data and transmission model as input. Details of the procedure are described in Supplementary note 2. Nodes associated with marked improvement stand as indicators of time points at which mutations predicted to have adaptive significance occurred. While we bring particular focus to the L-LCS^-^ treatment, identical analyses were performed on the remaining three treatments and are reported in Figure S8.

Figure 4 shows the result of the analysis for the L-LCS^-^ treatment. According to the maximum-*a-posteriori* procedure, genotypes that are ancestors of lineages where the probability of survival markedly improved are 2-I-A3 (generation 2, Ph I, lineage A3; marked A3 on the figure, +25%), as well as 1-I-F8 (+19%), 2-II-A5 (+1%), 3-I-A5 (+26%), 3-II-E2 (+1%), 4-I-E2 (+19%) and 5-I-E3 (+7%) and 5-I-C7 (+5%). Note that all these lineages had extant descendants at the end of the experiment. Moreover, 1-I-F8 and 2-I-A3 are ancestral to the entire population of lineages at generation 10.

**Figure 4.**
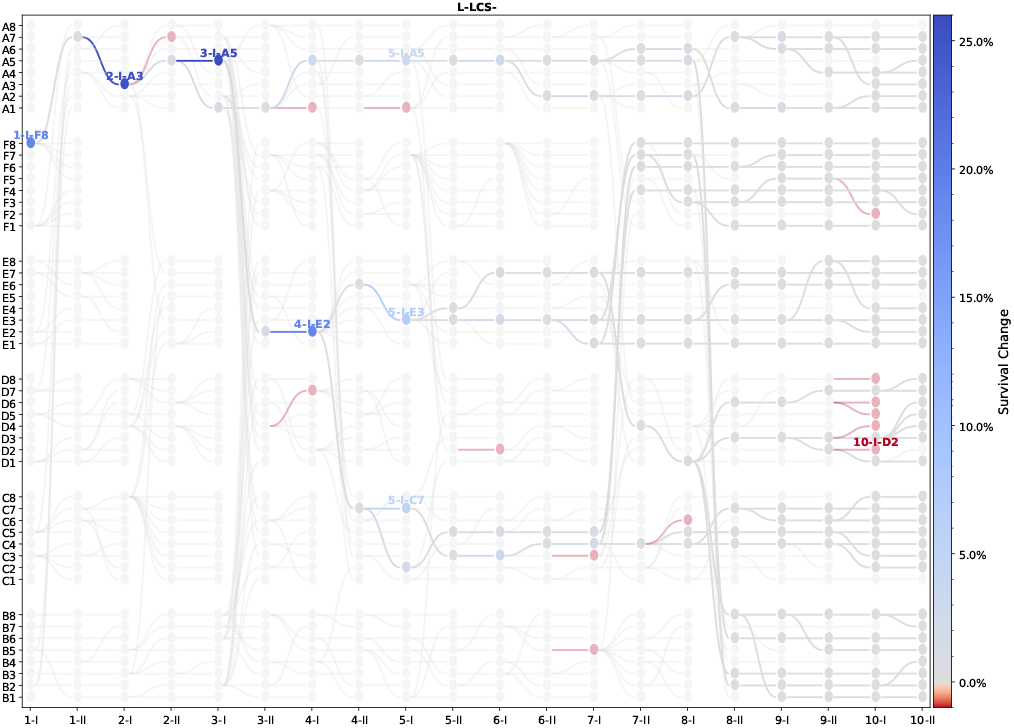
Nodes in the L-LCS^-^ genealogy predicted to mark adaptive change. Lineages are coloured according to predicted change in survival probability relative to the previous cycle. The belief-propagation algorithm used for analysis is described in Supplementary note 2. Here the model is beta with a grid of 100 points. Figure S8 shows this result for all four treatments.

In the next section, we describe how mutational data derived from whole genome sequencing – primarily of derived lines at generation 10 – aid elucidation of the genetic bases of fitness improvements.

### C. Propagating mutational data to the full genealogy

Whole genome sequence was obtained from all single WS (soma) genotypes derived from each extant lineage at the end of generation 10 (see Figure S13 for details), and from the single WS genotypes derived from the six selected microcosms marked in Figure 4. DNA sequence reads were aligned to the ancestral SBW25 genome leading to identification of a total of 9,271 mutations (across all four treatments). Because the 10 generations reported here were derived from LCS^+^, which had already experienced 10 prior generations of evolution, we removed those mutations (58 in total) that accrued during the previous experiment (See the full dataset in the “Data availability” section). While this represents an overwhelming number of mutations, in part, expected, because of the mutator status of the LCS^+^ genotype, mapping of mutations on genealogies is nonetheless possible. A belief-propagation algorithm similar to that described in the previous section was used. This builds upon the fact that an identical mutation found in two or more descendant lineages most likely emerged in a single ancestral lineage whose identity can be inferred from patterns of genealogical connections. The procedure is detailed in Supplementary note 3.

Given the large number of mutations, analysis began by firstly propagating the mutations from the end point (combined with additional information from the six focal lineages from L-LCS^-^) backward through each genealogy. The results, shown in Figure 5, reveal that lineages evolving under the S-LCS^-^ treatment (represented by dots) accumulated few mutations during the course of the experiment. This is as expected given that a single mutation underpins the transition between each phase of the life cycle: two mutations per cycle and thus ≈ 20 over the course of 10 generations. In contrast, lineages from both LCS^+^ treatments harboured ≈ 150 mutations, but some carried ≈ 250 mutations (represented by cross and plus symbols). Surprisingly, lineages from the L-LCS^-^ treatment (represented by diamond symbols) harboured, for some lineages, more than 350 mutations.

**Figure 5.**
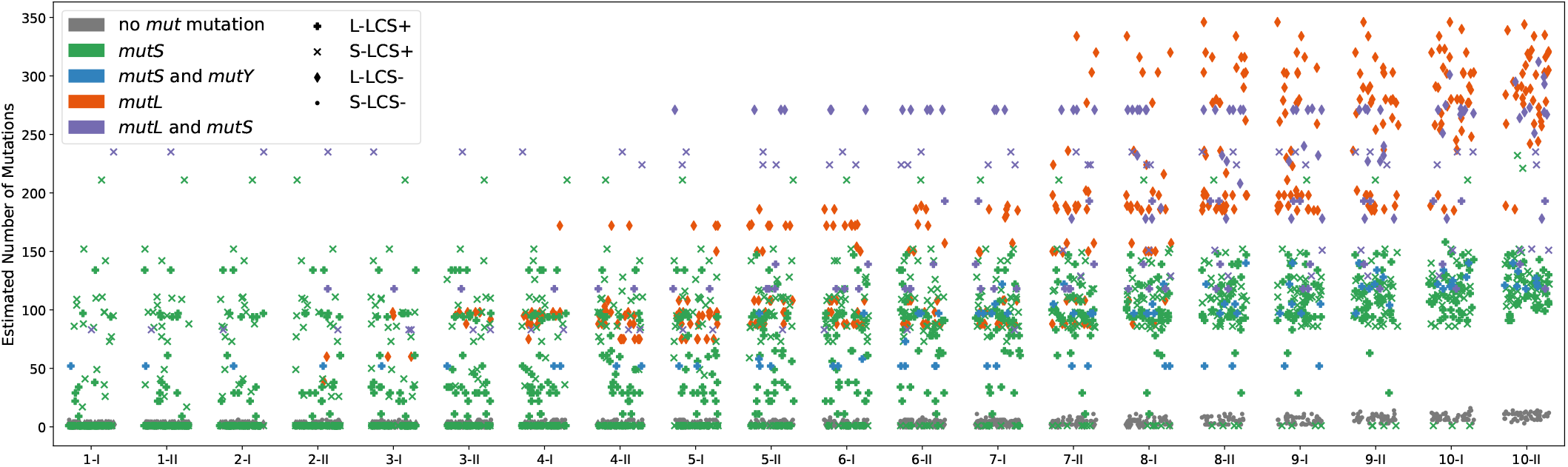
Reconstruction of the number of mutations through time. Each point is the number of new mutations (compared to the ancestor of the genealogy) predicted to be found in a single microcosm at a given phase in each cycle. The inference stems from propagating knowledge of mutations in the final derived genomes of extant lineages backward through the genealogy using the belief-propagation algorithm described in Supplementary note 3. Figure S10 shows the number of identified mutations without reconstruction through the belief-propagation regime. Figure S13 shows the number of mutations associated with each extent lineage (derived directly from genome sequence data).

Such large numbers of mutations are indicative of mutator genotypes. Mutation rate was not measured here, however mutation rates of bacteria harbouring mutations in primary components of the methyl-directed mismatch repair (MMR) system have been previously measured and shown to be elevated by between 70 and 300-fold (28, 29). In addition, for non-mutator lineages in a similar experimental setup, it is rare to fix more than a single mutation at each passage (29). From the number of mutations found in final clones, we can infer that on average 29.0 mutations per collective-generation are fixed for L-LCS^-^, and respectively 12.2 mutations/generation for L-LCS^+^, 13.0 mutations/generation for S-LCS^+^ and 1.0 mutation/generation for S-LCS^-^. Provided there are no changes in the probability of a mutation to fix, this estimate would indicate a 10 to 30-fold increase in mutation rate for mutator lines. That is, the same order of magnitude than reported in the literature.

While the presence of mutator genotypes was known for LCS^+^ lineages, we nonetheless interrogated all mutational data for evidence of MMR-defective types (*mutS, mutL, mutY* and *mutM*). The points in Figure 5, coloured according to genotype, show that microcosms harbouring lineages with large numbers of mutations also carry defects in mis-match repair genes (see Table 1). Again, while known for the *mutS*-based LCS^+^ lineages, these genotypes acquired additional mutations in MMR genes. For example, eight lineages in Block A of LCS^+^ evolving in large microcosms acquired a single non-synonymous change in the A/G specific adenine glycosylase *mutY* (G153D). Interestingly this *mutY* mutation did not generate more mutations (Figures 5 and S15), but is associated with a marked bias in the kind of mutations fixed (Figure S21).

The greatest number of mutations however were found in LCS^-^ lineages propagated in large microcosms. As more fully elaborated below, a single *mutL* (L103R) mutation (orange diamond symbols) arose early in the experiment, but seven lineages of block C also acquired an additional *mutS* (G707S) mutation (purple diamond symbols). Figure 6 and Figures S14 to S16 show these mutations in context of the genealogies apparent in each treatment.

**Figure 6.**
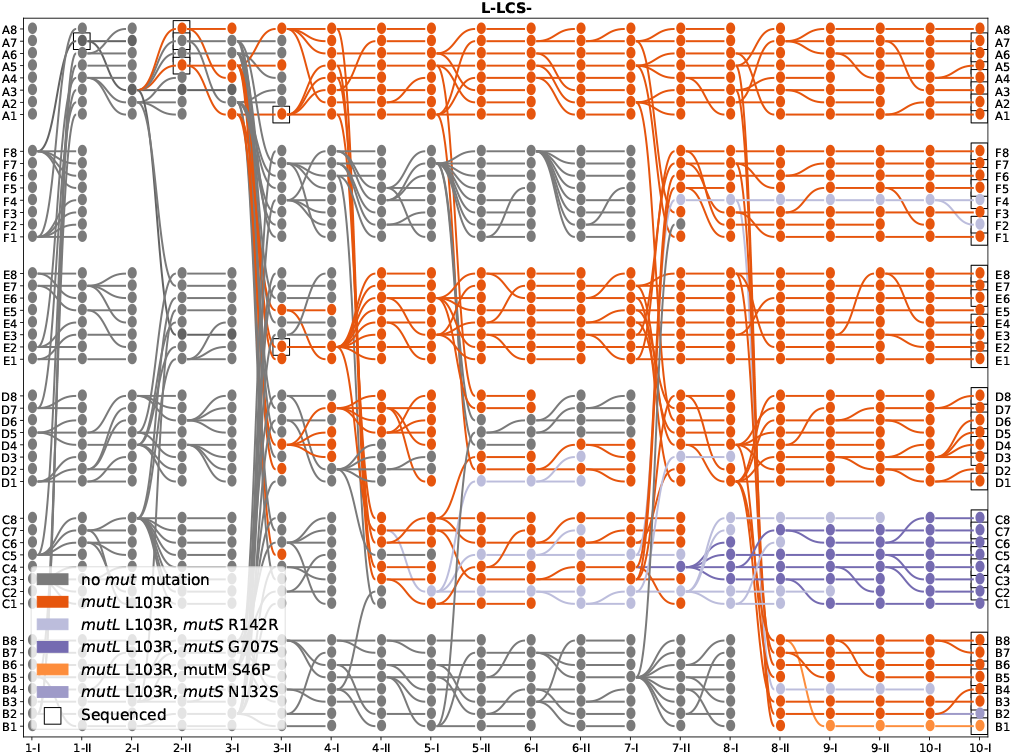
*mutL* and *mutS* mutations in the L-LCS^-^ genealogy. Lineages are colored according to *mutL* and *mutS* mutations. The mutation *mutL* L103R appeared in lineage 2-II-A5 and swept to fixation. Black squares identify sequenced lineages, with the belief-propagation algorithm (described in Supplementary note 3) allowing inferences to be extended to all nodes. Figures S14 to S17 show the same analysis for all four treatments.

As shown in Figure 4, the node marking greatest improvement in survival probability in L-LCS^-^ is in the vicinity of 2-I-A3 and 2-II-A5. Additional sequencing of these nodes showed that the point of origin of *mutL* (L103R) was immediately prior to 2-II-A5 (See Supplementary note 3-E and Figures S18 to S20 for details of the procedure and iterative use of the algorithm to incorporate new data).

### D. Experimental confirmation of adaptive changes

The combined information provided by the survival probability inference (Section B) and sequencing-information propagation (Section C), points toward an adaptive change having occurred in (or in the vicinity of) microcosm 2-II-A5. In this section, we describe additional experiments conducted to test this hypothesis.

Figure 7a-d shows the results of fitness assays performed around the adaptive points detected by Bayesian inference. These data, composed of measures of the probability to produce viable soma states at the end of Ph I (Figure 7a), probability to transition from soma to germ during Ph I (Figure 7b), probability to switch from germ to soma type during Ph II (Figure 7c), and overall life cycle fitness (Figure 7d), confirm a stepped increase in life cycle fitness of the focal lineage between generation two and four. Comparing the probabilities obtained through these additional experiments and the survival probabilities estimated by `colgen` shows agreement (Figure S9).

**Figure 7.**
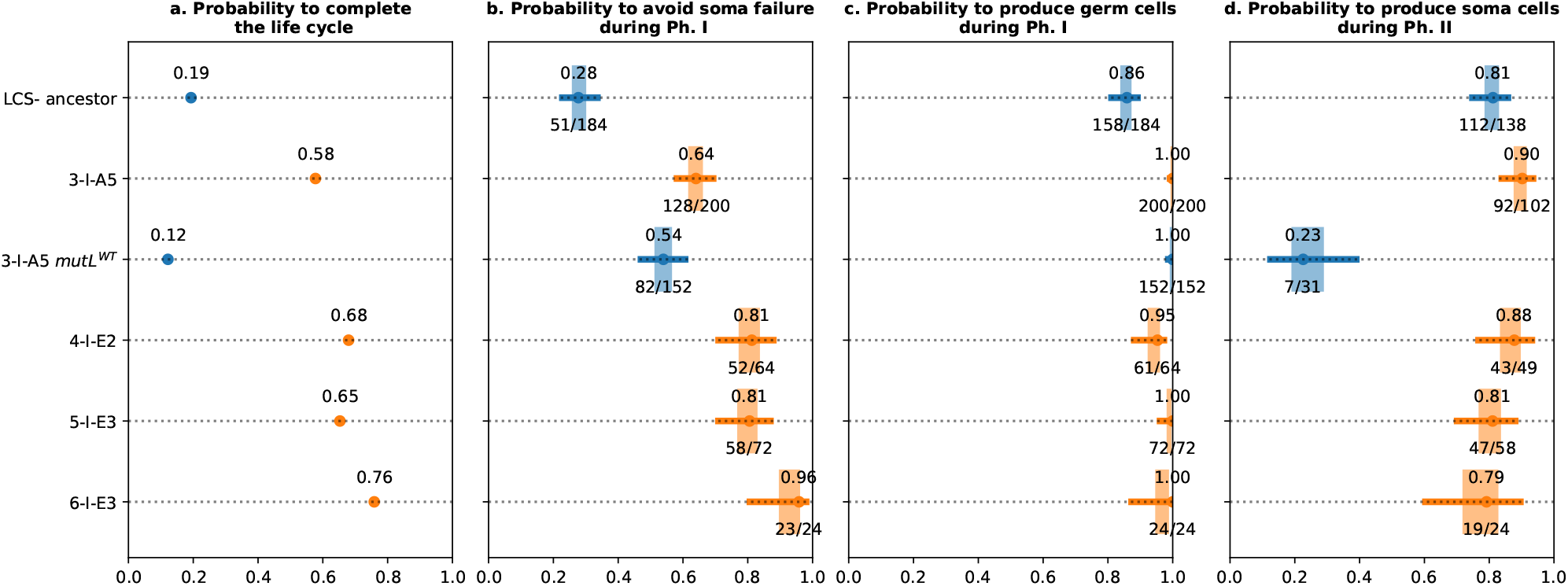
Probability and fitness estimates. Each line corresponds to a genotype derived from the ancestor LCS^-^, and grown in large microcosms. Points correspond to the maximum *a posteriori* value for the probability, the fraction of surviving over number of trials is written below the point. The box corresponds to the 50% Bayesian confidence interval and the whiskers correspond to the 95% Bayesian confidence interval (the method is described in Supplementary note 1). Orange depicts genotypes with the *mutL* L103R mutation; blue identifies genotypes without the *mutL* mutation.

Because prior analysis predicted a causal role for the *mutL* (L103R) mutation, this mutation (in genotype 2-II-A5) was reverted, via a single nucleotide change, to the ancestral state. The effects of this change are shown in Figure 7a-d. Data in Figure 7d demonstrate that at the level of overall life cycle performance, the *mutL* (L103R) mutation is the primary cause of enhanced life cycle success. Interestingly the effect is not significant for the two components of Ph I (Figure 7ab), but has a marked impact on ability of the germ stage to switch to the soma state. Further visual evidence for a seminal role of the *mutL* mutation in switching between life cyle states comes from observation of germ cells arising within soma directly on agar plates (Figure S22).

## Discussion

The evolution of multicellular life continues to receive attention from both empiricists and theoreticians (8, 30–39). In this work, using the experimental *Pseudomonas* system, we have explored the efficacy of a previously derived genetic switch controlling life cycle transitions, used knowledge of genealogical connections among nascent multicellular entities to predict adaptive events, combined these data with DNA sequence information and, for one particular mutation, demonstrated a causal connection to improved fitness.

The genetic switch, that in a prior experiment had delivered adaptive benefit to competing lineages through significant improvement in control of expression of soma- and germ-like phases, proved, in this work, central to subsequent evolutionary success. This held for continued selection in standard microcosms, but also for selection in large vessels. LCS^+^ lineages suffered few extinctions during further propagation in standard microcosms, whereas persistence of LCS^-^ lineages in the same diameter microcosms proved challenging, with frequent death events caused by failures at all three life cycle phases.

Both LCS^+^ and LCS^-^ lineages struggled initially to find solutions to producing enduring mats in large diameter microcosms, but in both instances this capacity showed improvement during the course of the experiment. Notable in the L-LCS^-^ treatment, was the early discovery of a mutation in *mutL* that effectively converted LCS^-^ to LCS^+^. In the absence of this mutation – and thus marked improvement in capacity to transition between phases of the life cycle – all lineages in the L-LCS^-^ treatment would have gone extinct by the third generation (evident by excluding reproduction events in Figure 3 and following the fate of lineages to the first extinction). Thus, mutator-fuelled switching increased reliability of the transition between life cycle phases, but the elevated mutation rate likely also played a role in discovery of genetic changes that led to the formation of mats not only capable of colonising the surface of large diameter microcosms, but with sufficient structure to ensure persistence over the six days of Ph I.

The prevalence of mutators associated with successful lineages was particularly apparent for lineages evolving in large microcosms and shows that the selective regime strongly favours mutational-based genetic switches. These switches deliver benefit through subsequent rounds of selection, despite presumed costs arising from the inevitable load of deleterious mutations. However, there are several reasons to suspect that these costs maybe less significant than in standard mutation accumulation experiments. One important factor is the fact that the bottleneck is *selective* with only cells expressing the correct phenotype transitioning to the next phase (40). Such selective bottlenecks are likely to purge some deleterious mutations, thus reducing the mutational burden usually associated with hypermutagenesis (41). Additionally, the six and three-day periods of Ph I and II, respectively, provide ample time for both purifying selection and compensatory evolution. Finally, collective level selection stands to counteract accumulation of deleterious mutational effects: should a lineage carrying a costly mutation pass through the bottleneck, such that it is unable to produce the next phenotype of the cycle, then its failure is compensated by replacement from the pool of extant lineages. Note that mutations that reduce growth rate, and thus appear maladaptive on an individual cell basis, can prove adaptive at the collective level (13) and over longer time scales (42).

While L-LCS^+^ lineages were founded by a genotype carrying a *mutS* allele, a further *mutY* G153D mutation arose early in a single Block A lineage and went to fixation by generation 10. That the mutation swept, suggests that it may have contributed to further improvements survival probabilityof collectives (Figure S8), however, it is not inconceivable that *mutY* G153D hitchhiked with a mutation that improved performance of the soma phase. Interestingly this mutation did not add to the overall number of mutations (Figures 5 and S15), but it did introduce a marked bias in the kinds of mutation found in these lineages (Figure S21), with recent work suggesting that the ensuing transversion-biased mutational spectrum might afford greater possibility for discovery of adaptive mutations (43). Interrogation of sequence data for changes in repeat-tract length in genes with predicted DGC or PDE activity did not identify obvious candidate effectors of phenotype switching (as previously observed in WspR (17) see Figure 1).

In large microcosms founded by the LCS^-^ lineage, in addition to the early and key *mutL* mutation, a further *mutS* G707S mutation arose in Block C microcosms. As with the *mutY* allele arising in the *mutS* background, the *mutS* G707S mutation, although arising late in the experiment (at generation seven), fixed (in Block C) within a further two generations (Figure S14). While the adaptive significance of this mutation was not further investigated, microcosm 8-I-C4 was identified by `colgen` as a point that marked significant improvement in survival probability (Figure 4). Moreover, multilocus mutators have been shown to provide additional benefits to populations evolving under strong selection (44).

The fact that no additional mutators evolved in either the S-LCS^-^ or the S-LCS^+^ treatment suggests that additional opportunities for mutator alleles to hitchhike with beneficial mutations were provided in large micrcocosms. This makes intuitive sense given the combination of selective challenge, where founding genotypes were initially poorly adapted, a three-fold increase in population size (compared to standard microcosms), and the fact that transitions through the life cycle require passage through single cell bottlenecks. Under such conditions there are likely numerous beneficial mutations upon which mutator alleles can hitchhike, moreover, selection is expected to favour cells that switch phenotype early because descendants of such cells have a heightened change of passage through the bottleneck.

A notable difference in this study compared to previous work (17, 18) is mode of analysis, particularly that delivered by `colgen`. The first and arguably most valuable component of `colgen` is the simple visual representation of evolutionary dynamics via which evolutionary patterns previously invisible are revealed, allowing, for example, contributions of chance, history and selection to be directly observed. This is further aided by comparisons among the fates of lineages founded by LCS^+^ and LCS^-^ genotypes. The former being endowed with the adaptive Muts-WspR based switch – the product of prior evolutionary history – whereas the latter, by virtue of reversion of the *mutS* allele to wild type, has this component of history removed.

LCS^+^ lineages propagated in standard microcosms suffered few extinction events (Figure 3a), which can be directly attributed to prior evolutionary history and is consistent with adaptive value of the switch. Notable was the worsening of performance after generation three Figure S2. This likely reflects the downside of few initial extinction events, resulting in a shift in the relative contributions of selection acting between- versus within-lineages (45). In the absence of lineage-level death-birth events, selection operates primarily within lineages leading to erosion of traits favouring success of lineages. But as lineage-level death-birth events occur, selection operates both within and between lineages, with, presumably, some equilibrium level ultimately being achieved.

In contrast, for LCS^-^ lineages propagated in the same standard microcosms, multiple extinction events occurred (Figure 3C). Beyond providing further evidence of the adaptive value of the switch, extinction and concomitant reproduction events are indicative of the hand of chance – both good and bad luck. In fact, as there is no overall improvement in the frequency of extinction events in S-LCS^-^ lineages – and none detected by `colgen` (Figure S8), it is likely that the observed dynamics are driven almost entirely by chance.

Chance is an inherent factor in all evolutionary change, but a major factor affecting extinction events (Figure 3). Chance underpins ability to transition (by mutation) between life cycle phases, with mutational switches greatly improving the likelihood that transitions are successfully achieved, but chance also plays a role in mat durability with subtle differences determining whether a mat endures the required six-day period. Additionally, chance contributes to whether cells composing the next life cycle phase reach a frequency sufficient to ensure detection.

Extinction, while terminating evolution of affected lineages, provides opportunity for those that are extant to reproduce, and here again, chance is important. Death – often a consequence of bad luck – means that extant lineages, which may be equally susceptible to extinction, serendipitously gain further opportunity to explore genetic (and thus phenotypic) space, and through such exploration, the possibility presents that these lineages, at some future time point, might discover fitness-enhancing mutations: persistence, and its counterpartbserendipity, is everything (24).

A single lineage from Block B in the S-LCS^-^ treatment is a case in point (Figure 3c). The lineage in microcosm b3 produced offspring after the first generation, at generation three, and at generation four, but all failed soon after. Nonetheless, by virtue of persistence at generation five, where the seven competing lineages failed, this lineage went on to leave eight offspring at generation six; by end of the experiment, this single generation-five survivor had given rise to eleven extant offspring. Without further study, the extent to which spread of this lineage reflects adaptive change is unclear, but it is plausible that success to this point was largely a matter of chance. This is also reflected in life cycle fitness data in Figure S4 where there is no compelling evidence for the hand of selection, with the fitness of several extant lineages at end of the experiment, being equal or worse than the ancestral type.

A further factor relevant to considerations that recognise the value of sheer persistence, stems from the fact that lineage-level selection, as implemented in the experiments described here, involves a “death-birth” dynamic. This is in contrast to a “birth-death” process in which offspring replace extant collectives, regardless of fitness. While game theory approaches show that death-first dynamics favour the evolution of cooperation (46, 47), in the context of this experiment a birth-death process would likely diminish the value of persistence.

Also notable in the dynamics of S-LCS^-^ lineages (but also evident elsewhere) are abrupt extinctions of all lineages (within single blocks) following several generations of successful transitions (Figure 3c). See for example Blocks F and C at Ph I of generation six. While possibly just chance, this more likely reflects deleterious effects of prior mutational history that due to lineages having traversed various mutational paths during prior cycles have exhausted possibilities for realisation of mutations critical for the next life cycle transition.

Turning to LCS^+^ and LCS^-^ lineages propagated in large microcosms, similar patterns manifest, however, also apparent are dynamics consistent with the hand of selection. LCS^+^ lineages, although replete with genetic switch (a product of past history), suffered high levels of extinction during the first few generations, primarily due to premature failure of the soma stage, with such events becoming increasingly rare from generation five. While chance was dominant early on, and the effect of prior history minimal (as expected given that the founding genotypes had not previously experienced selection in large microcosms), sweeps arising from individual lineages, likely underpinned by adaptive change, resulting in fixation of successful types by generation 10.

LCS^-^ lineages propagated in large microcosms, as described above, narrowly escape wholesale extinction. Having eliminated prior history through elimination of the genetic switch, the fate of these lineages were determined by both the requirement to find solutions to switching and construction of robust mats. Again, as mentioned, a key adaptive change – a mutation in *mutL* – occurred at generation two, with this lineage becoming the progenitor of all extant lineages (across all blocks) at generation 10. A combination of chance and selection were sufficient to overcome – and more than compensate for – prior evolutionary history.

Representation of genealogies via `colgen`, while invaluable in this work is generally application to numerous kinds of lineage selection experiments where mixing of lineages is avoided, as in the “propagule-pool” (as opposed to “migrant-pool” or “mixed”) mode of reproduction (18, 29, 48). This holds irrespective of whether collectives are clonal (17) or communities (14, 49, 50).

Beyond providing graphic visualisation, `colgen`, by virtue of the underlying Bayesian model, has utility in allowing mutational data obtained from the end of the experiment to be propagated backward throughout the genealogy. This allowed sense to be made of the hundreds of mutations present in mutator lineages, which are typically overwhelmingly difficult to deal with. But in addition, amongst the noise of mutations, `colgen` proved invaluable at identifying microcosms likely to contain adaptive mutations. Detailed investigations into L-LCS^-^ led to identification of the *mutL* mutation that was pivotal to the success of descendant lineages.

Graphical probabilistic models (such as Bayesian networks) for statistical inference have been used in haplotyping of pedigree (51), gene network inference (52) and genetic diseases risk assessment (53). Bayesian network analysis presents several advantages. Firstly, it is designed to work with partial data: inference can be performed even when sequence data is missing or partial. Secondly, such modes of analyses allow seamless integration of new data, with the possibility of adding new sequencing or fitness information to refine inferences. Thus, Bayesian analysis makes possible iterative protocols that alternate between inferences based on existing data and targeted new experimentation. Finally, the statistical model of observation and transmission of characters is relatively simple to define and can be adjusted for different purposes. While the models used above were minimal in their assumptions, it is possible to conceive and develop more complex mechanistic models as required.

It is evident that life cycles based on mutation can deliver benefits to nascent multicellular types. However, life cycles in extant multicellular organisms are typically effected via developmental mechanisms requiring changes in patterns of gene regulation. While it would be interesting to continue propagation of lineages containing mutation-dependent switches to observe future refinement, there is minimal possibility that true developmental control could be achieved using an experimental protocol that requires soma and germ phases to express distinct morphologies on agar plates. This has led to a revision in which this requirement has been removed. Indeed, selection under a revised protocol that remains unchanged except for the fact that the germ-line phase is assessed by dispersal (and not colony morphology) has shown that lineages capable of transitioning between life cycle phases by developmental control are eminently achievable (Summers *et al*. forthcoming).

## ACKNOWLEDGEMENTS

GD acknowledges the John Templeton Foundation (#62220); GD and PBR acknowledge funding from *Origines et Conditions d’apparition de la vie* (ANR-10-IDEX-001-02). PBR is grateful to the Max Planck Society for generous core support and funding from the Marsden Fund Council from government funding administered by the Royal Society of New Zealand. PBR thanks J. Carlos Hernandez-Beltran and Kaumudi Hassan Prabhakara for comments on a draft of the manuscript. We thank Mathieu Forget for contributions while an intern at the NZIAS.

## AUTHOR CONTRIBUTIONS

PBR, experimental design and conception; PR and DR, conduct of experiment; GD development and implementation of `colgen`; all authors analysed data and wrote the MS.

## DATA AVAILABILITY

All the data used in this manuscript (genealogy records, mutation identification from read alignments, fitness assays) and the code necessary to reproduce all the figures from this dataset is available in the Zenodo repository: https://doi.org/10.5281/zenodo.11170875. `colgen` v2.0b1 used in this manuscript is also available on Zenodo: https://doi.org/10.5281/zenodo.7342768. The sequencing data files will be available within the European Nucleotide Archive (accession number pending).

## Supplementary Information

**Figure S1.**
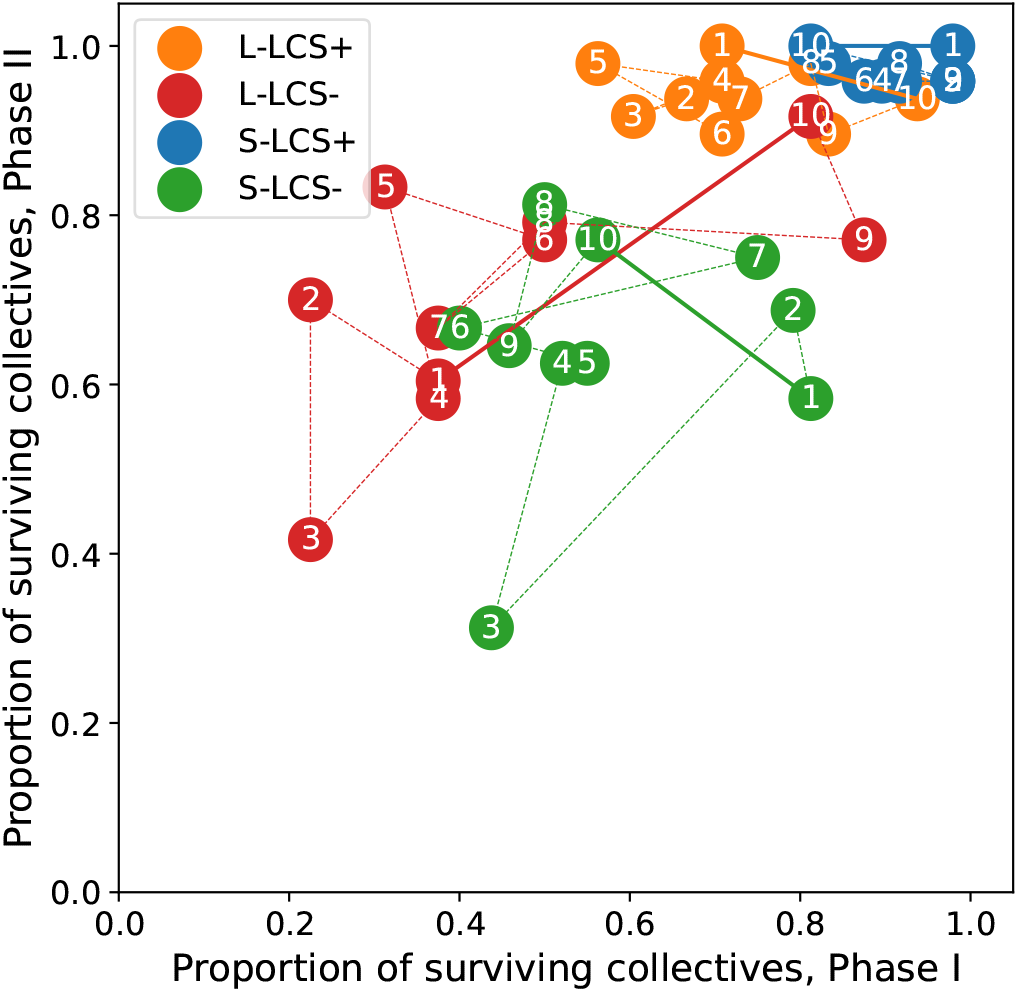
Proportion of surviving lineages in Ph I and Ph II for each cycle. Each circle corresponds to the proportion of surviving lineages for the cycle that is labelled. Each color is a different treatment, LCS^+^: Lineage 17 from (1) with a life cycle switch, LCS-: same lineage with switching capacity eliminated, S: standard microcosms, L: large microcosms. The dashed lines link consecutive cycles, the thick lines link the first and last cycle, representing the resulting change over the whole experiment. Note that for all treatment but L-LCS-, there exists a trade-off between survival in the two phases.

## Supplementary Note 1: Fitness assays

Previously, measures of fitness have involved lineage-level competition experiments against ancestral types that are often difficult to interpret and challenging to conduct. Observation of the genealogies of lineages as shown in Figure 3 led to realisation that the probability of avoiding extinction is a good predictor of future evolutionary success. Henceforth, measures of fitness were determined, not by competition with an ancestral type, but by measuring the number of lineage extinction events after replicate lineages of single test genotypes have gone through a further life-cycle generation.

New experiments were conducted with the same protocol as for the main lineage-selection experiment and performed for a single cycle, each time starting with a population of eight microcosms founded by the same genotype. At the end of phase I, the presence of a mat was recorded and the density of SM cells was measured though plating. At the end of phase II, the density of WS cells was estimated through plating.

This experiment was conducted on average 44 times for each of the 48 genotype and microcosm size combination tested (hence a total of 2120 microcosms). For each genotype, we estimated the probability of three events: *M* the production a mat at the end of phase I, *S*_*I*_ the production of cells with the switched phenotype SM cells at the end of Ph I, *S*_*II*_ the production cells with the switched phenotype WS at the end of Ph II. Further, we defined the “fitness” of a genotype as its probability to survive a full two-phase cycle, as the probability to survive both phases, hence *F* = **P**(*M* ∧*S*_*I*_ ∧*S*_*II*_).

We estimated the three probabilities by maximum of likelihood on a beta-binomial model, with a non-informative prior (uniform in [0,1]). As a consequence, ff there were *n* independent trails and *k* successes, the estimate of probability is simply the proportion of successes,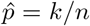 and the Bayesian confidence interval is given by the quantiles of a beta distribution parametrised as Beta(1 + *k*, 1 + *n*−*k*).

**Figure S2.**
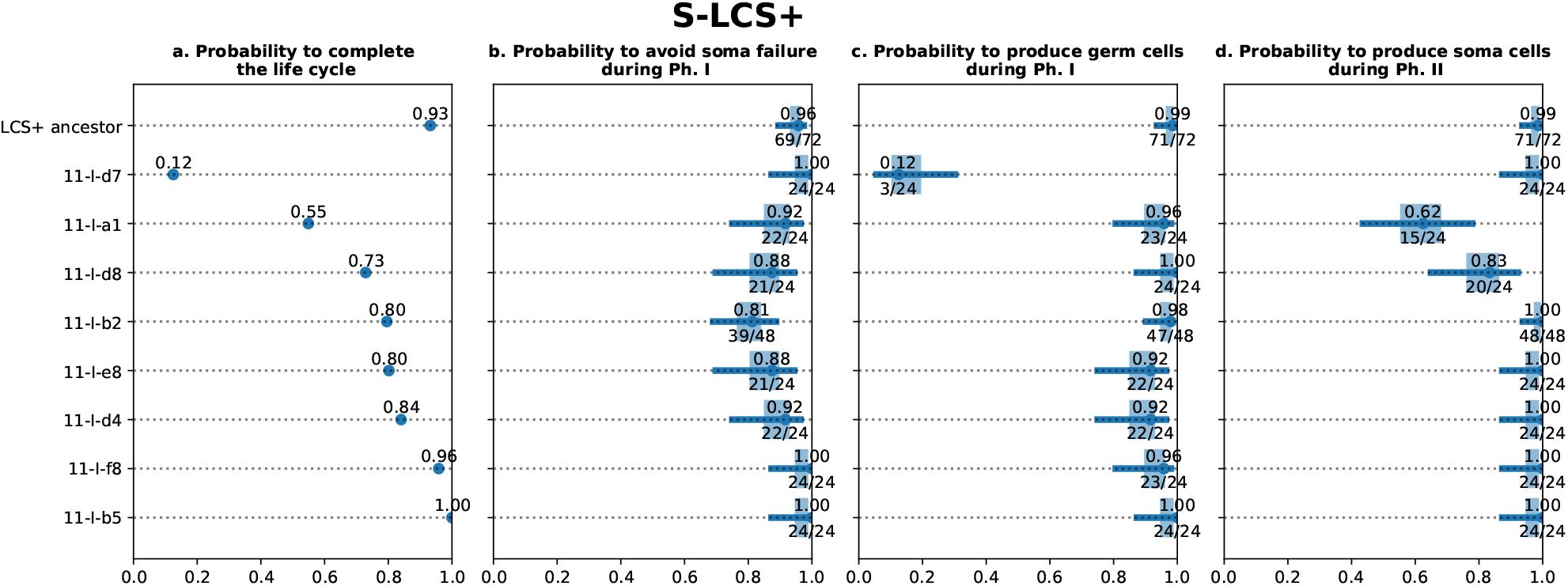
Probabilities in S-LCS^+^ endpoints. Each line corresponds to a genotype derived from the ancestor LCS^+^, and grown in small microcosms. Same caption than Figure 7.

**Figure S3.**
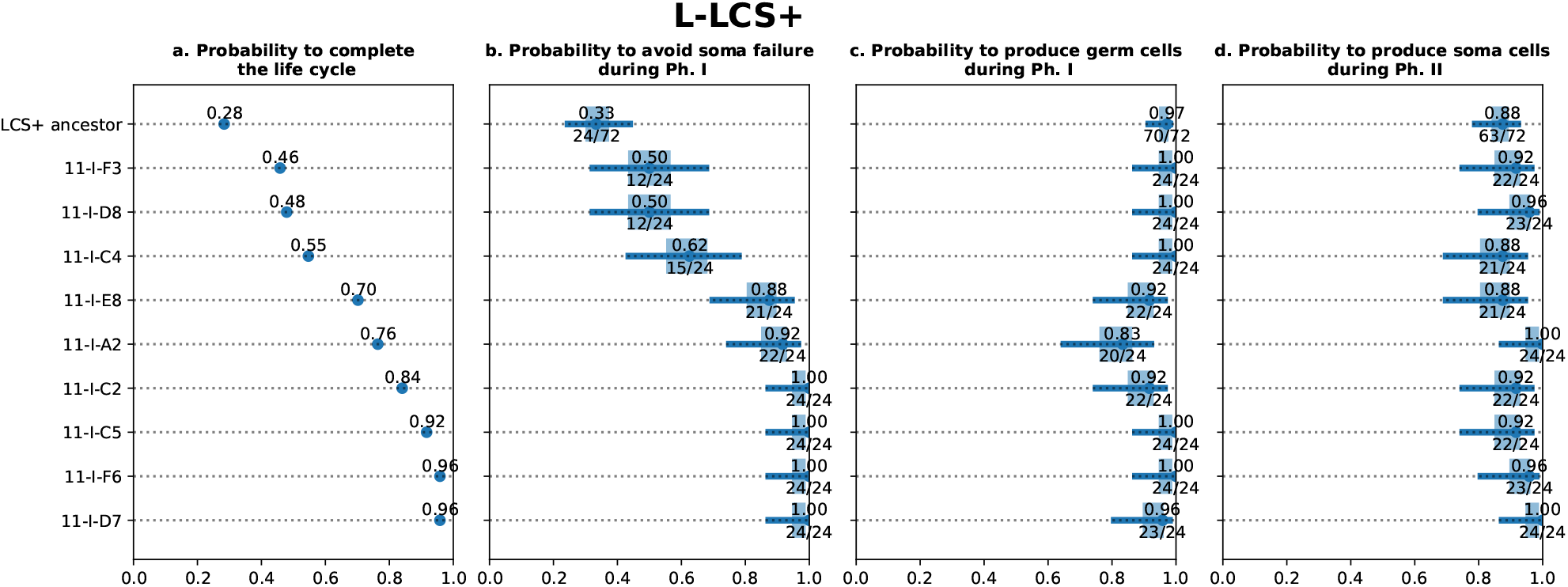
Probabilities in L-LCS^+^ endpoints. Each line corresponds to a genotype derived from the ancestor LCS^+^, and grown in large microcosms. Same caption than Figure 7.

**Figure S4.**
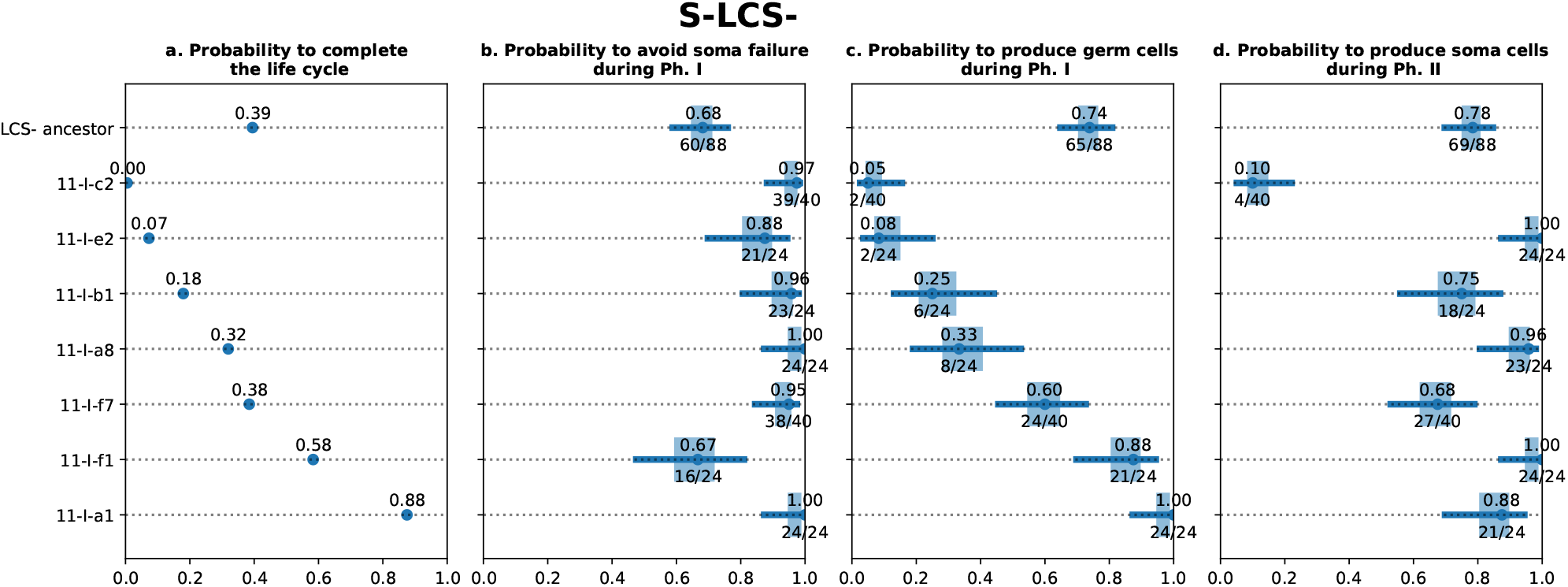
Probabilities in S-LCS^-^ endpoints. Each line corresponds to a genotype derived from the ancestor LCS^-^, and grown in small microcosms. Same caption than Figure 7.

**Figure S5.**
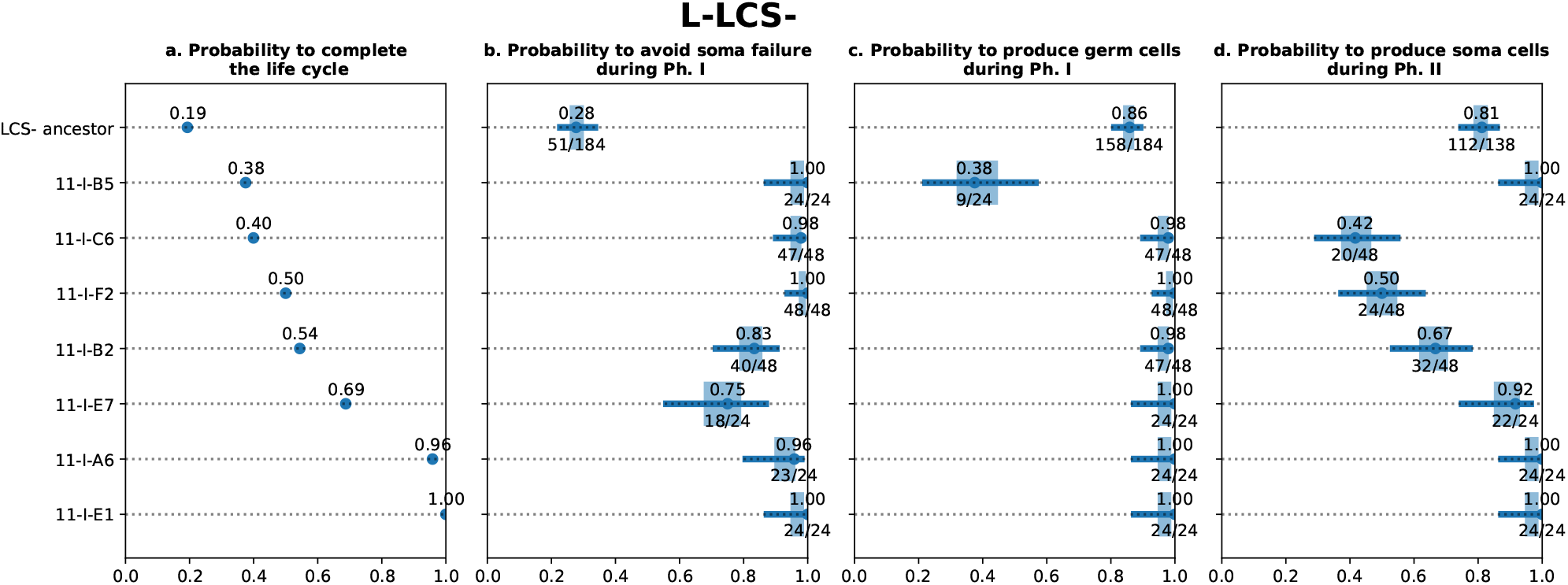
Probabilities in L-LCS-endpoints. Each line corresponds to a genotype derived from the ancestor LCS^-^, and grown in large microcosms. Same caption than Figure 7.

## Supplementary Note 2: Adaptive value estimation

We describe a method to assign to each collective within the genealogy a most probable adaptive value. In technical terms, the “most probable adaptive value” refers to a maximum likelihood estimation (MLE) that is a way to assign, under a defined graphical probabilistic model, a value to a parameter that maximise the probability of the observed result. The adaptive value is here a hidden variable: one does not simply observe the adaptive value of a collective, but performs experiments that provide information on its true value.

This endeavour requires first encoding of both the observations and the hidden characteristics of the collectives in a model, that takes the form of a joint probability distribution. Second, a method to find the most likely assignation of an adaptive value to each culture within the genealogy, for a given mutation rate, will be presented. Conversely, parameter inference allows assignment of the most likely value of a mutation rate for a given configuration. Finally, the iterative method of Expectation-Maximisation will be used to concurrently fit to the data both the adaptive value and the mutation rate.

### A. Model structure

**Figure S6.**
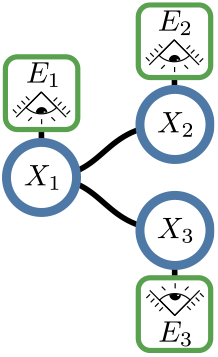
Bayesian Network. Bayesian network corresponding to a genealogy of three collectives. Collective 1 has two offsprings: collectives 2 and 3. Vertices are random variables, edges are conditional dependence. The hidden variables *X* encode the survival probability of the collectives, while the observed variables *E* encode the outcome of the experiment (survived or not).

Each collective genealogy 𝒯 = (*V, ε*) is a forest containing *N* numbered vertices: *V* = 1…*N* which are the collectives, and an edge linking each collective *i* to its parent *p*_*i*_: ε = (*i, p*_*i*_)_*i*=1…*N*_.

The model assigns to each collective an observed and a hidden variable. For a collective *i*, let *E*_*i*_ be the random variable encoding whether this collective went extinct (*E*_*i*_ = 1) or not (*E*_*i*_ = 0). In the field of probabilistic graphical models the observed values such as the (*E*_*i*_)_*i*=1…*N*_ are called “evidence”.

The first assumption of the model is that there exists a hidden random variable for each collective *i*, called *X*_*i*_, representing its survival probability. In probabilistic terms, this means that the conditional probability of extinction of a collective *i* knowing the value of the survival probability *x*_*i*_ is exactly the value of *x*_*i*_:

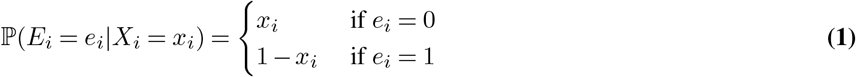

The second assumption of the model is that the survival probability of a collective is transmitted from one collective generation to the next, with an unbiased small variation. In probabilistic terms it means that if the parent collective has survival probability 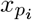, then the survival probability of its offspring collective follows a distribution with mean 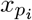 Several probability distributions families can be used. Colgen implement two of them: a beta distribution and a “jump” distribution family parameterized by a single scalar *σ*.

For the beta distribution:

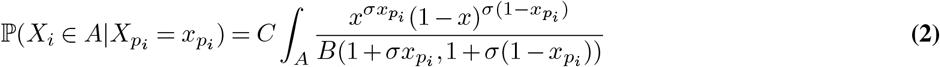

With 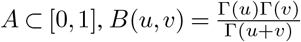, Γ the Gamma function, and *C* a scaling factor ensuring that 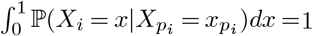

For the Jump distribution:

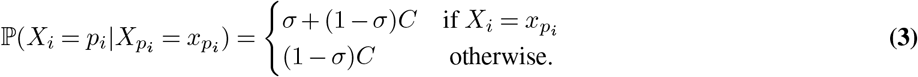

With *x*_*i*_, 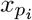 both in (1*/C*, 2*/C*…*k/C*, 1), and *C* a scaling factor ensuring that 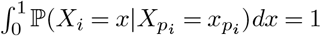

During the experiment, the values of *E*_*i*_ = *e*_*i*_ (extinct or not) are observed, whereas the values of *X*_*i*_ are “hidden”.

### B. Bayesian inference

Consider a collective *i* with associated survival probability *X*_*i*_ and experimental outcome *E*_*i*_. The prior probability distribution of *X*_*i*_ is given by Equation 2. Consider for an instant that 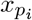, the adaptive value of its parent is a fixed parameter. Bayes’ Theorem gives the expression of the probability of *X*_*i*_, conditional on the observed value of *E*_*i*_:

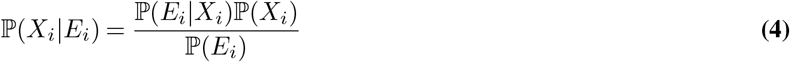

Where ℙ (*X*_*i*_) is the prior probability of *X*_*i*_, ℙ (*E*_*i*_|*X*_*i*_) is the observation model from Equation 1 and ℙ (*E*_*i*_) is the probability of the evidence. The maximum likelihood estimate (or more precisely the maximum a posteriori estimate, MAP) of the value of *X*_*i*_ is given by taking the value of *x* that maximises ℙ (*X*_*i*_ = *x*|*E*_*i*_), note that the value of the MAP does not depend on ℙ (*E*_*i*_), which is a simple scaling constant. Observing the outcome of the experiment (survival or extinction) has an effect on the posterior: the whole distribution (and consequently the MAP) is shifted toward higher values if the collective survived, or lower values if the collective went extinct.

The problem lies, as often in Bayesian statistics, in the prior probability of *X*_*i*_ (2). Indeed, in the context of the experiment, this probability depends on the state of all the ancestors of the collective *i*. Applying Bayes theorem to all those interdependent observations quickly becomes a complex undertaking without a proper method. Evidence propagation, presented hereafter, is this optimal method.

The next section details the technical steps required to find the most probable values of *X*_*i*_ throughout the genealogy.

### C. Finding the most likely configuration of the genealogy

The configuration of the full genealogy is the knowledge of the state of all vertices. Hence, it is encoded as a joint distribution that maps each configuration to a probability:

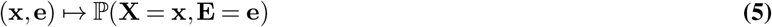

Where bold letters are used to represent vectors of dimension *N*, **x** = (*x*_*i*_)_*i*∈*V*_, **X** = (*X*_*i*_)_*i*∈*V*_, **E** = (*E*_*i*_)_*i*∈*V*_, **e** = (*e*_*i*_)_*i*∈*V*_. The likelihood of a given set of adaptive values **x** is defined as the probability of observing this value, conditional on the state of all collectives:

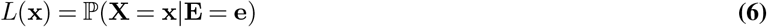

The maximum a posteriori configuration of the network is given by finding the value of **x** that maximises *L*(**x**). The naïve, brute force method to find this value is to compute the *L* for all Card(*ω*)^*N*^ possible configurations (where Card(*ω*) is the number of possible value of *X*. If the adaptive value is discretised in a hundred of values, this gives 100^48^ computations (in the order of the number of atom of earth ≈ 10^50^, and way too much even for modern computers).

However, the probability distribution can be efficiently factorised thanks to the definition of conditional probabilities whose structure is represented in the graph: each pair of nodes not linked by an edge is conditionally independent of one another. As an example consider the small network in Figure S6. The factorisation will go like this:

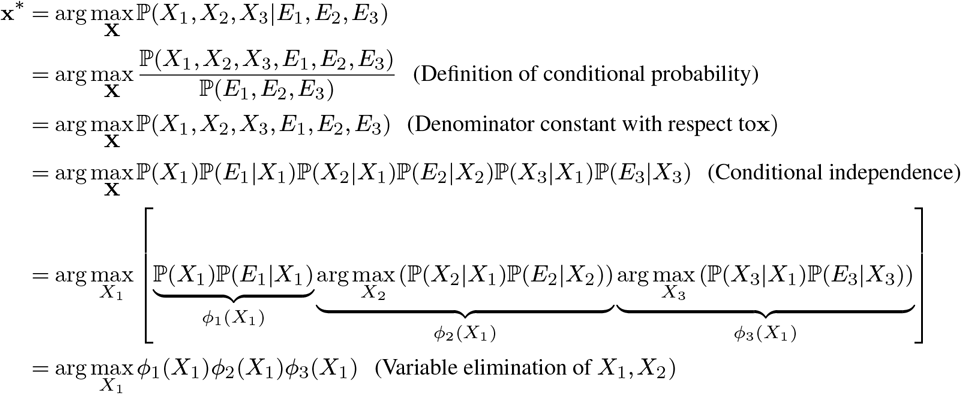

Additionally, the “maximum” operation is distributive, which allows for efficient computation of local maxima (*ϕ*_2_, *ϕ*_3_, called messages) that depend on a single variable. This constitutes the essence of the Max-Product Belief propagation algorithm ((3), Chapter 13).

### D. Expectation-Maximisation procedure

Now, how can one find the value of the parameters of the model ? The Expectation-Maximisation (EM) algorithm (4), which might be one of the most influential advances in modern computational statistics and is widely used in Gaussian Mixtures, and Hidden Markov Models, provides an iterative solution.

First, note that for any given configuration of the network (values of **X** and **E**), finding the maximum likelihood estimate of the parameter (mutation rate *σ*) is a simple matter of computing the value of the parameter that maximises the likelihood ℙ (**X, E**)).

Now, starting from an arbitrary value of parameters (say,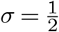), one may compute the Maximum A Posteriori configuration of the network, that assigns a value to each hidden variable. This constitutes the Expectation (E) step. Then, a new value of the parameter is computed using the Maximum of Likelihood estimate. This constitutes the maximisation step (M). The procedure can be iterated until convergence. This algorithm is guaranteed to reach a local maximum in likelihood, but may not reach the global maximum. However, its conceptual simplicity means that it is often the best heuristic available for practical applications.

### E. Pre-processing specific to the present life-cycle experiment

The mat-selection experiment described in the main text features two alternating phases (selection for germ, selection for soma) involving very different processes and are likely to yield different survival probabilities. As a consequence, two sub-trees were extracted for each genealogy, one for Ph I and the other for Ph II (Figure S7) and analysed separately.

**Figure S7.**
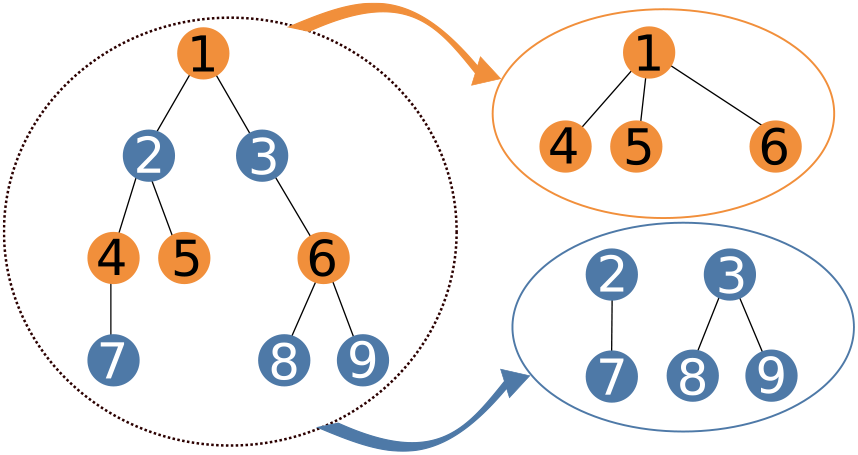
Subtree extraction procedure. Phase 1 (selection for a somatic mat and germ-like cells) and phase 2 (selection for the reversion to soma-like cells) are qualitatively different and are treated separately. In order to achieve this, each collective genealogy is split into a phase 1 and a phase 2 tree that are analysed on their own, and re-assembled for visualisation.

---

**Figure S8.**
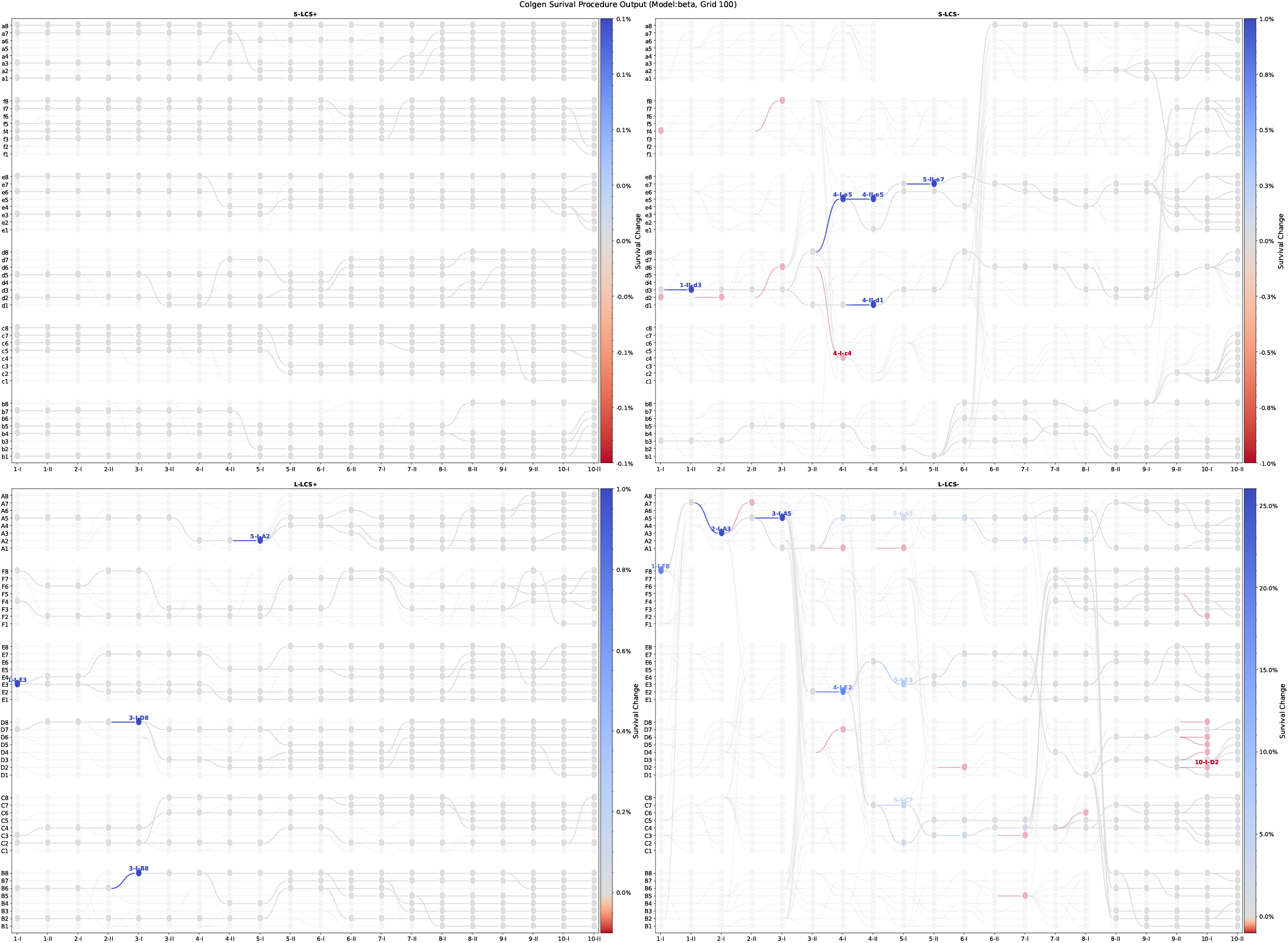
Nodes in the genealogies predicted to mark adaptive change. Model Beta, Grid:100, Sigma-max:1500.

**Figure S9.**
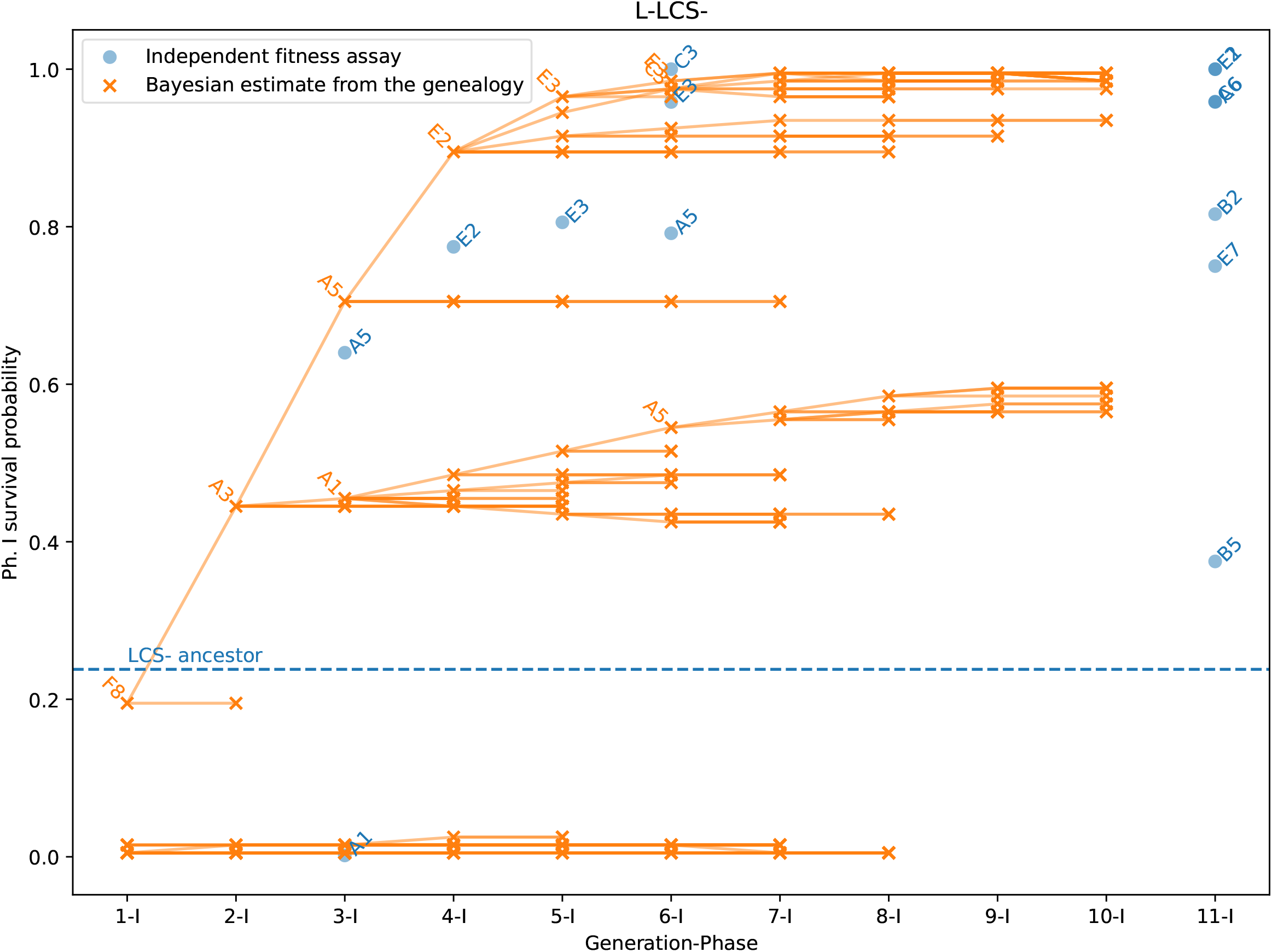
Phase I survival probability as estimated by independent fitness assays (blue, data from Figures 7 and S5) and the Bayesian beta model of Colgen (Orange, Sigma=1500, Grid=100, from Figure 4).

## Supplementary Note 3: Sequencing and mutation history reconstruction

### A. Number of mutation identified by direct sequencing

Figure S10 represents the number of mutations identified in each sequenced lineage. Note that the L-LCS^-^ has a significantly higher number of mutations than S-LCS^-^ at cycle 10 depsite being started with the same genotype. It seems that it has became a hypermutator line.

**Figure S10.**
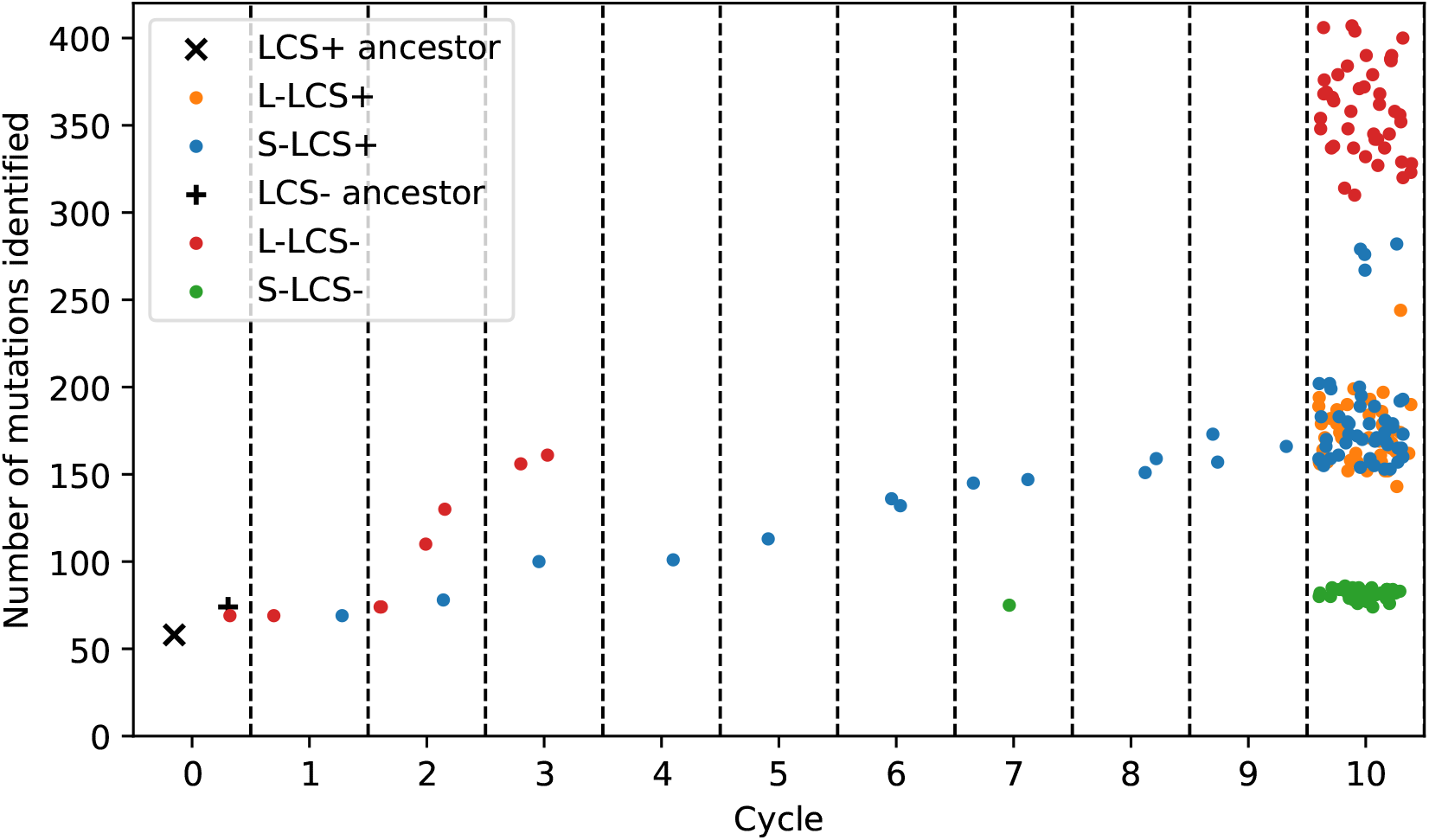
Number of mutations identified by direct sequencing. Each circle is a sequenced culture. Mutations are identified by mapping shorts read onto the reference SBW25 genome (GCF000009225_2). See materials and methods for details. The x-axis correspond to the cycle, the x-position is randomized within each cycle to avoid overlap. Compare this figure with Figure 5 (Note that in Figure 5, the mutations present in the ancestor are filtered out) and Figure S11 that show this data propagated through the whole genealogy.

**Figure S11.**
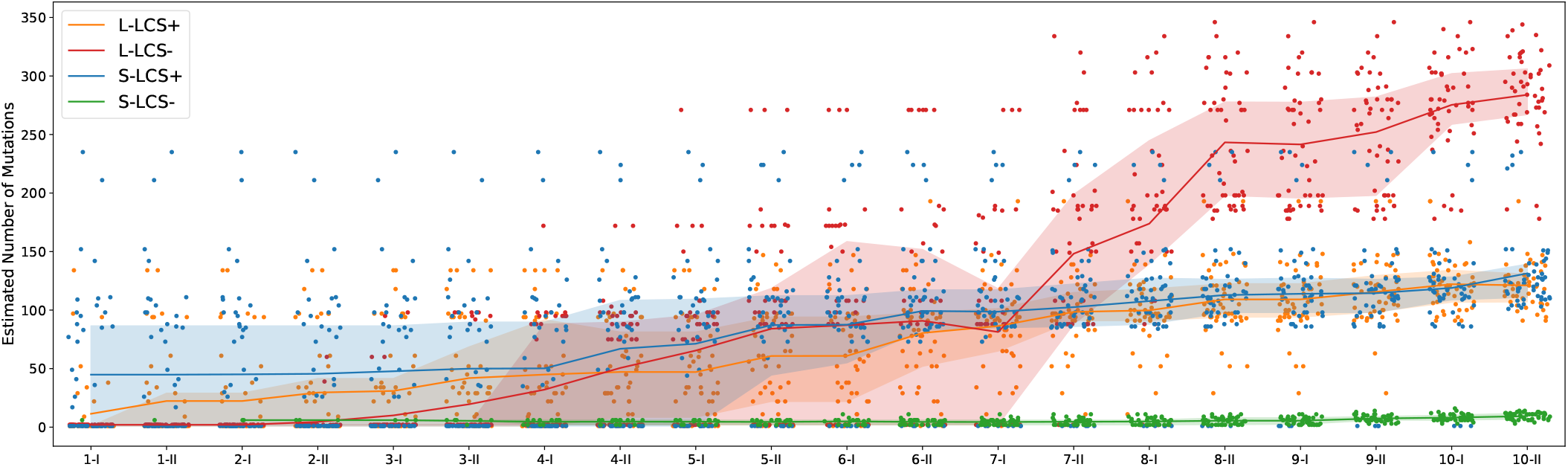
Number of new mutations reconstructed by belief propagation. Each circle is a sequenced WS genotype. The x-axis corresponds to generation of the life cycle, the x-position is randomized within each cycle to avoid overlap. Compare this figure with Figure S10 that shows these data only for mutations identified by sequencing (and includes mutations present in the ancestor). Lines correspond to the average number of mutations per treatment, shaded areas correspond to 50%-quantils.

### B. Propagation of mutational data from sequencing through genealogies

We use the belief propagation algorithm presented in Supplementary note 2, but the hidden variable *X*_*i*_ is now the presence (*X*_*i*_ = 1) or absence (*X*_*i*_ = 0) of the mutation within the lineage. The observed variable *E*_*i*_ is replaced by a conditionally Bernoulli variable *O*_*i*_ encoding the event “the mutation of interest was identified in sequences obtained from collective *i*” for the genotype that was sequenced. The observation model must be specified:

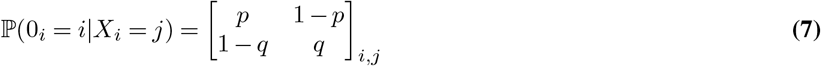

Where *p* and *q* represent, respectively, the probability of true negative and true positive reporting. In the following *p*=*q* =1. Finally, the heredity model is specified as:

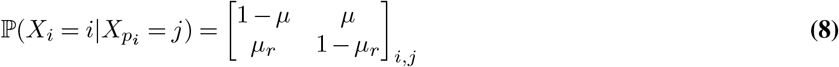

Where *µ* is the probability that the mutation appears within one cycle and *µ*_*r*_ the probability of losing the mutation. In the main text, arbitrary values of *µ* = 0.001 and *µ*_*r*_ = 0.00001 have been selected.

### C. Number of mutations

Figures S11 and S13 represent the number of mutations predicted in each node by the belief propagation algorithm.

### D. Mut mutations

We focused on the mutations in genes known to encode components of the methyl-directed mismatch repair system (*mut* family: *mutS, mutL, mutY* and *mutM*, 17 mutations were identified, see Table 1). The results are shown on the genealogies in Figures S14 to S17.

**Table 1.** Description of the mutations in the methyl-directed mismatch repair system identified in the dataset. Figure S12 shows in which lineages they were identified.

| Gene | Position | Mutation | Annotation | Description |
| --- | --- | --- | --- | --- |
| *mutY* ← | 355688 | C→T | G153D (GGC→GAC) | A/G-specific adenine glycosylase |
| *mutL* → | 586966 | C→T | S5L (TCG→TTG) | DNA mismatch repair protein |
| *mutL* → | 587050 | A→G | E33G (GAG→GGG) | DNA mismatch repair protein |
| *mutL* → | 587260 | T→G | L103R (CTG→CGG) | DNA mismatch repair protein |
| *mutL* → | 587555 | C→T | L201L (CTC→CTT) | DNA mismatch repair protein |
| *mutL* → | 587737 | A→G | N262S (AAC→AGC) | DNA mismatch repair protein |
| *mutL* → | 588126 | (G)5→6 | coding (1174/1902 nt) | DNA mismatch repair protein |
| *mutL* → | 588551 | C→T | A533A (GCC→GCT) | DNA mismatch repair protein |
| *mutS* ← | 1300356 | C→T | G707S (GGC→AGC) | DNA mismatch repair protein |
| *mutS* ← | 1300673 | +A | coding (1802/2592 nt) | DNA mismatch repair protein |
| *mutS* ← | 1300783 | A→G | R564R (CGT→CGC) | DNA mismatch repair protein |
| *mutS* ← | 1300986 | T→G | T497P (ACC→CCC) | DNA mismatch repair protein |
| *mutS* ← | 1301392 | A→G | R361R (CGT→CGC) | DNA mismatch repair protein |
| *mutS* ← | 1302049 | A→G | R142R (CGT→CGC) | DNA mismatch repair protein |
| *mutS* ← | 1302080 | T→C | N132S (AAC→AGC) | DNA mismatch repair protein |
| *mutS* ← / ← PFLU_1165 | 1302576 | G→A | intergenic (-102/+68) | DNA mismatch repair protein/putative membrane protein |
| *mutM* ← | 6343361 | A→G | S46P (TCC→CCC) | formamidopyrimidine-DNA glycosylase |

**Figure S12.**
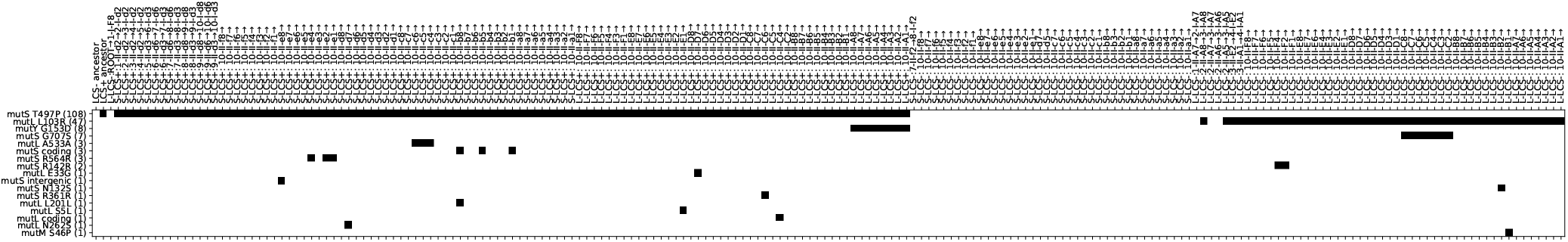
Identified mutations of the methyl-directed mismatch repair system in the sequenced genomes. A black square indicate that the mutation was identified. See Table 1 for a description of the mutations.

**Figure S13.**
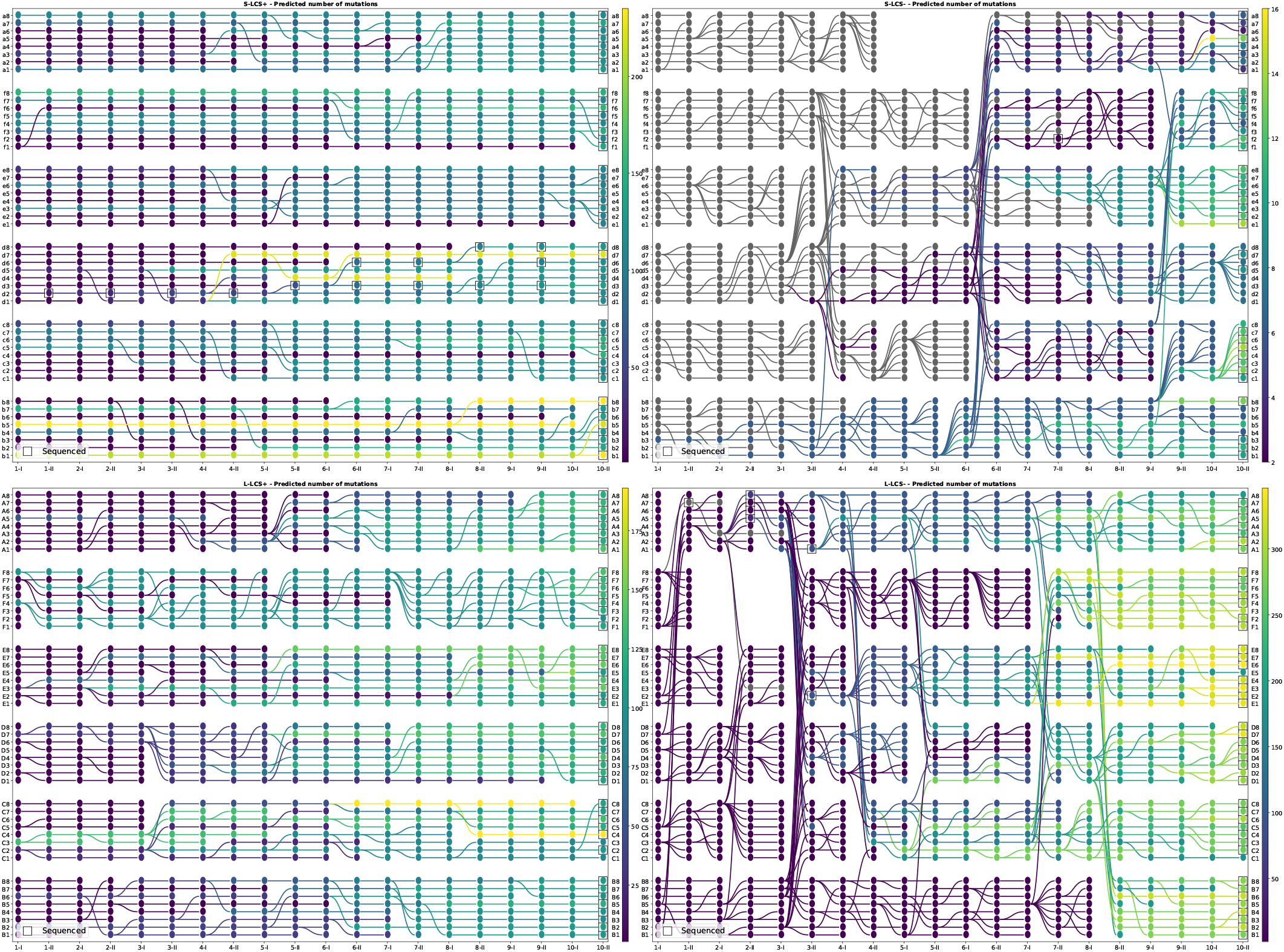
Estimated number of mutations per node. Sequenced genotypes are marked with a black square.

### E. Finding the location of the L103R *mutL* mutation

Figures S18 to S20 explain how we pinpoint the location of L103R *mutL*.

When we only had the endpoints of L-LCS^-^ sequenced, the L103R *mutL* mutation was in all the sequenced nodes, so the best prediction that could be done was “in the last common ancestor” See Figure S18.

However, the information obtained through the survival estimation algorithm prompted further investigation around 2-II-A5 and 3-II-E2, so additional sequencing was conducted of nodes 1-II-A7, 2-II-A5, 2-II-A6, 2-II-A7, 2-II-A8, 3-II-E2 and 3-II-A1. The mutation is present in 2-II-A5, 2-II-A8, 3-II-E2 and 3-II-A1 and absent in the other nodes (Figure S19).

The propagation procedure of `colgen` output two origins of mutL (Figure S20), in 2-II-A5 and 2-II-A8, because it is highly unlikely that it was in 1-I-A3 and was not transmitted to 2-II-A6 and 2-II-A7.

This mean that the mutation probably appeared between the single cell bottleneck at the end of 1-II-A7 and the single cell bottleneck at 2-II-A5 and 2-II-A8. However, not all cells within 1-I-A3 carry the mutation.

### F. Mutation bias in *mutY*

Figure S21 shows the mutation biais in lineages with a *mutY* mutation.

**Figure S14.**
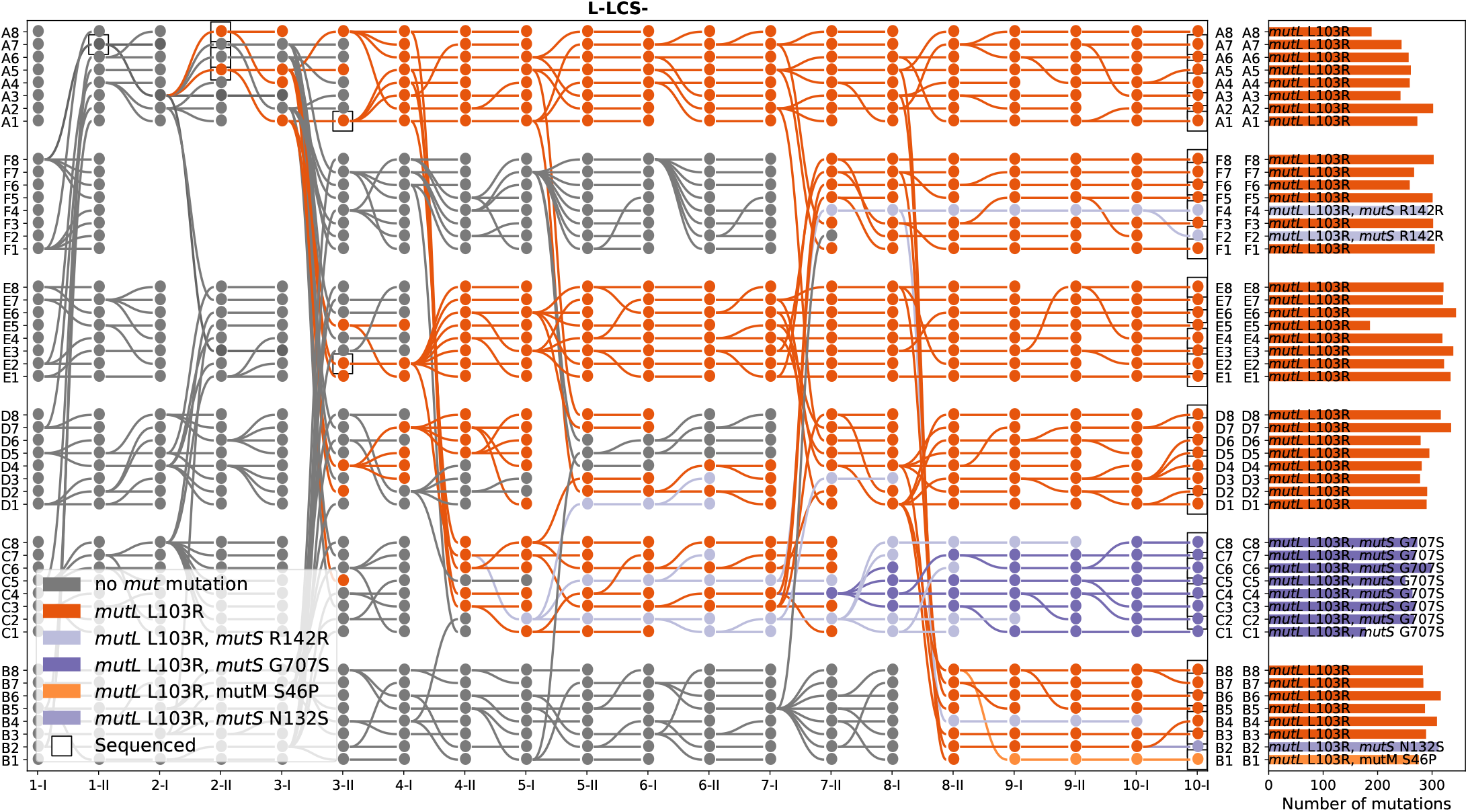
Mut mutations and number of mutations at the endpoint for L-LCS^-^.

**Figure S15.**
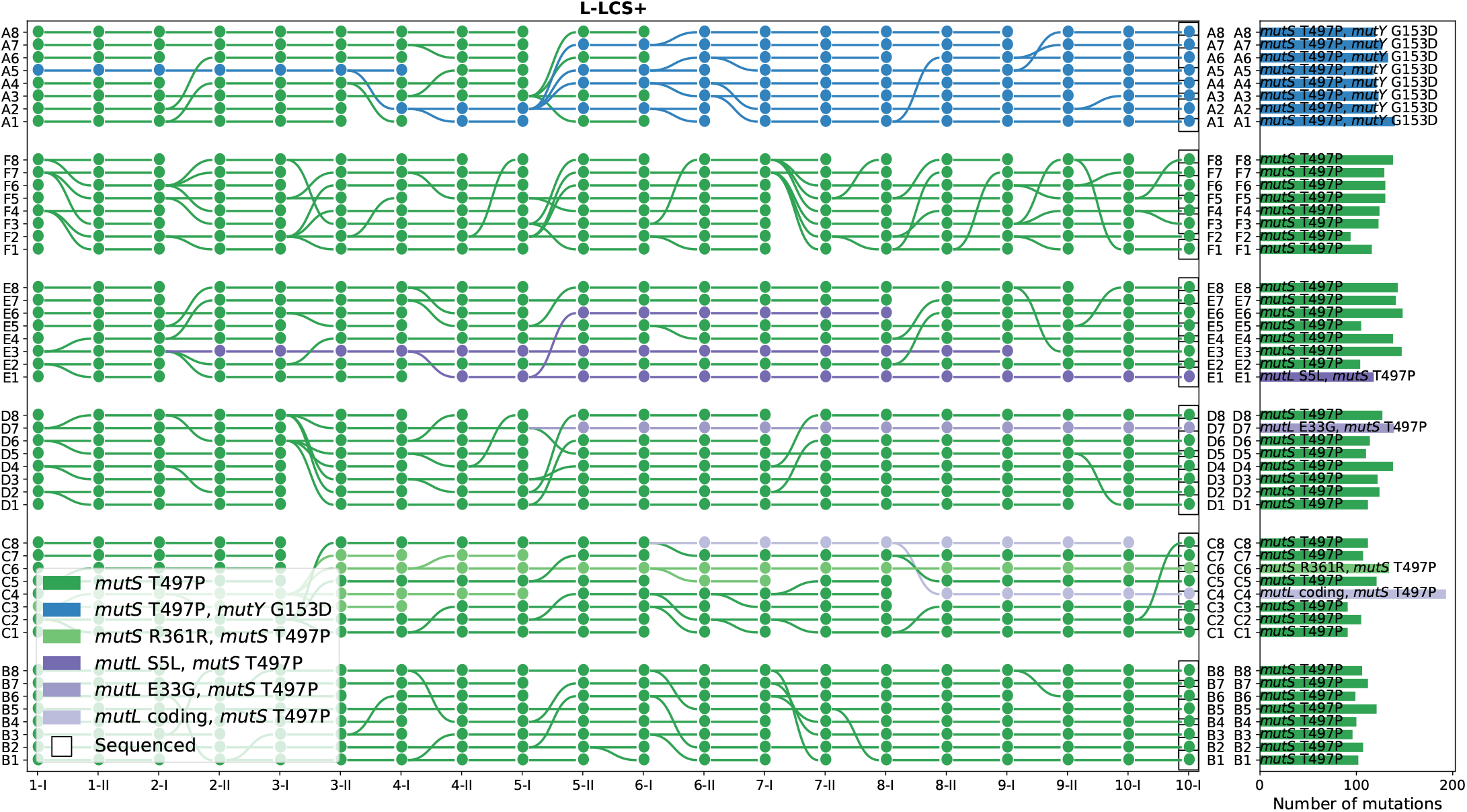
Mut mutations and number of mutations at the endpoint for L-LCS^-^.

**Figure S16.**
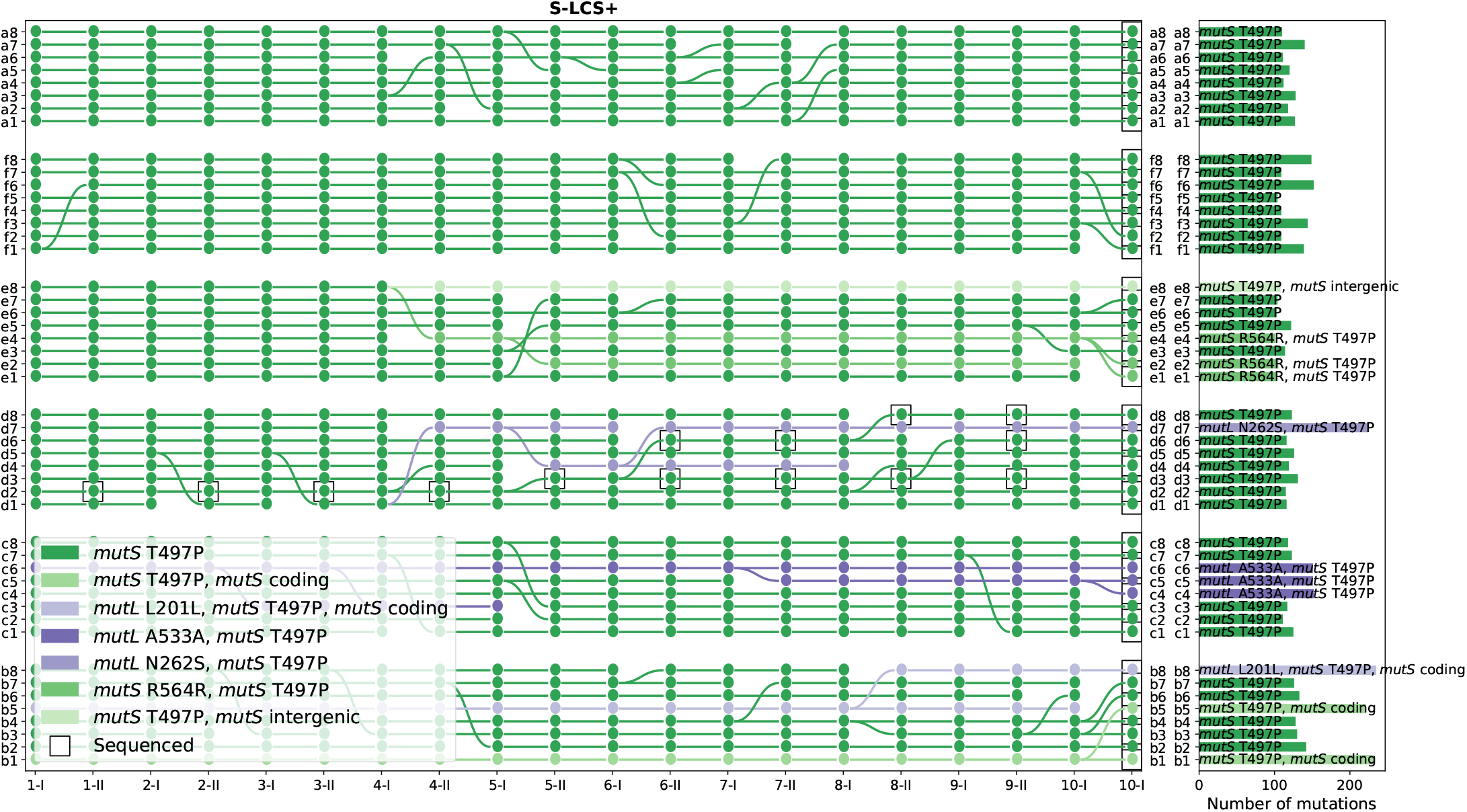
Mut mutations and number of mutations at the endpoint for S-LCS^-^.

**Figure S17.**
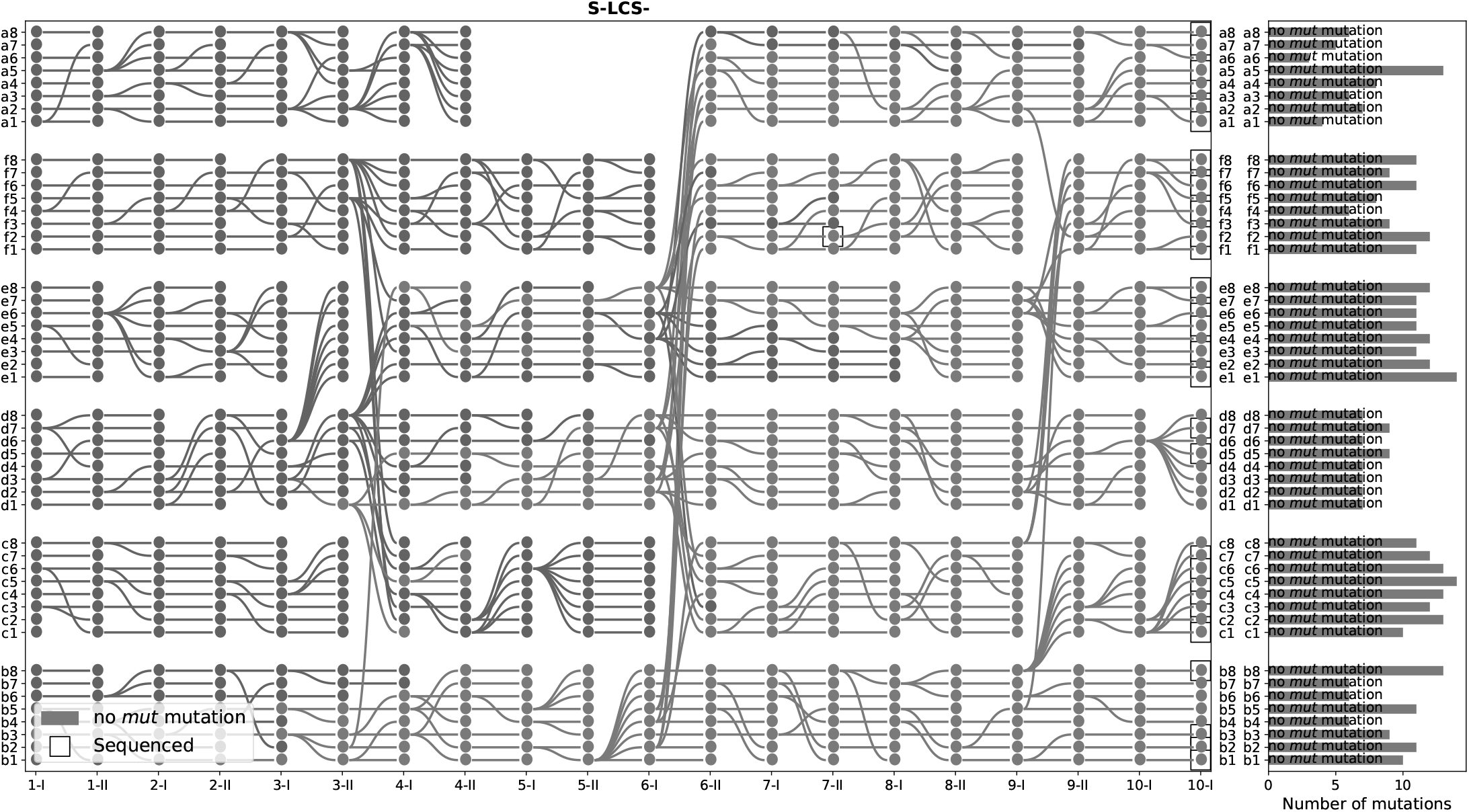
Mut mutations and number of mutations at the endpoint for S-LCS^-^.

**Figure S18.**
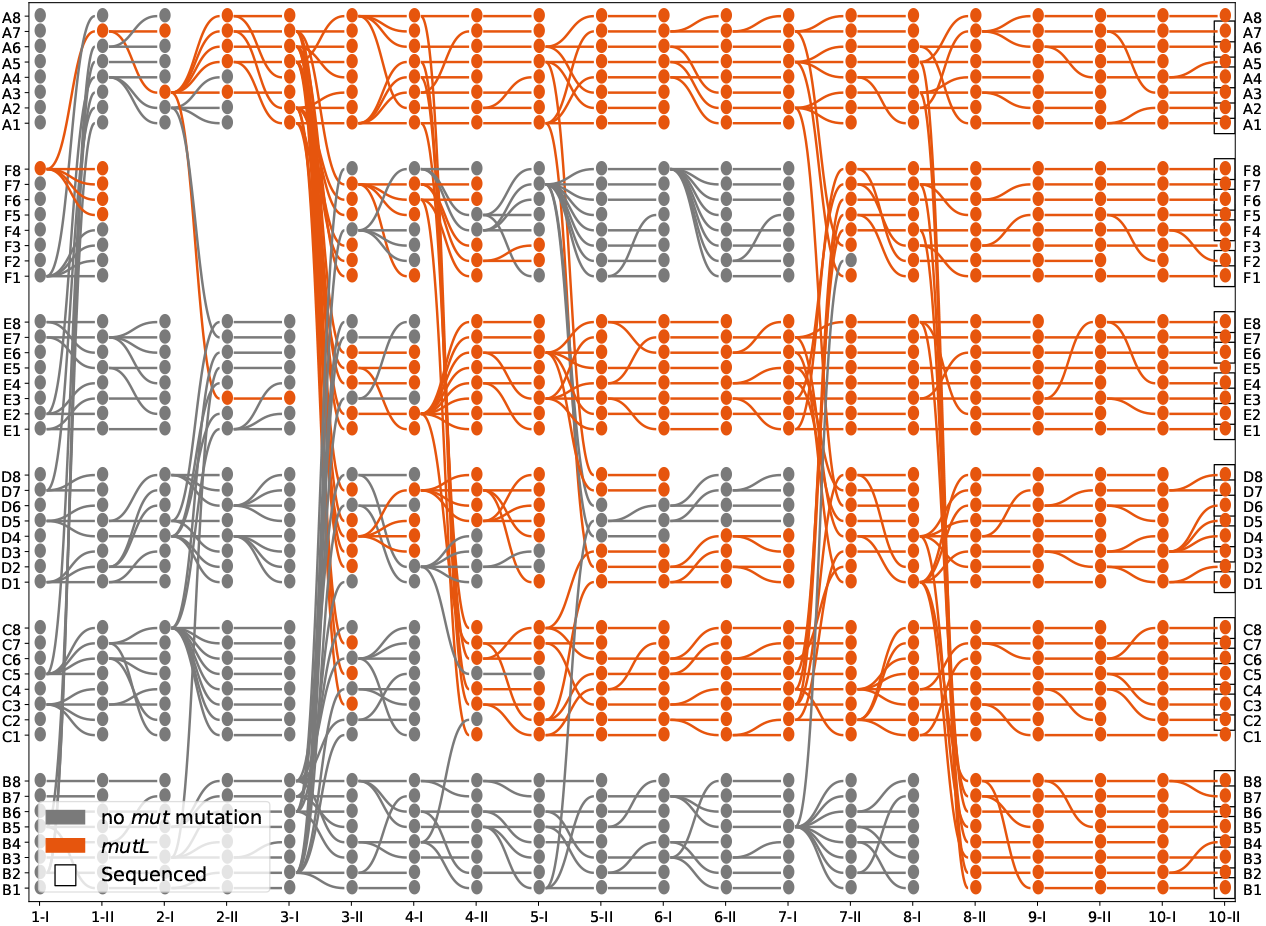
Origin of the *MutL* L103R mutation, estimated using only the sequencing data from the endpoints.

**Figure S19.**
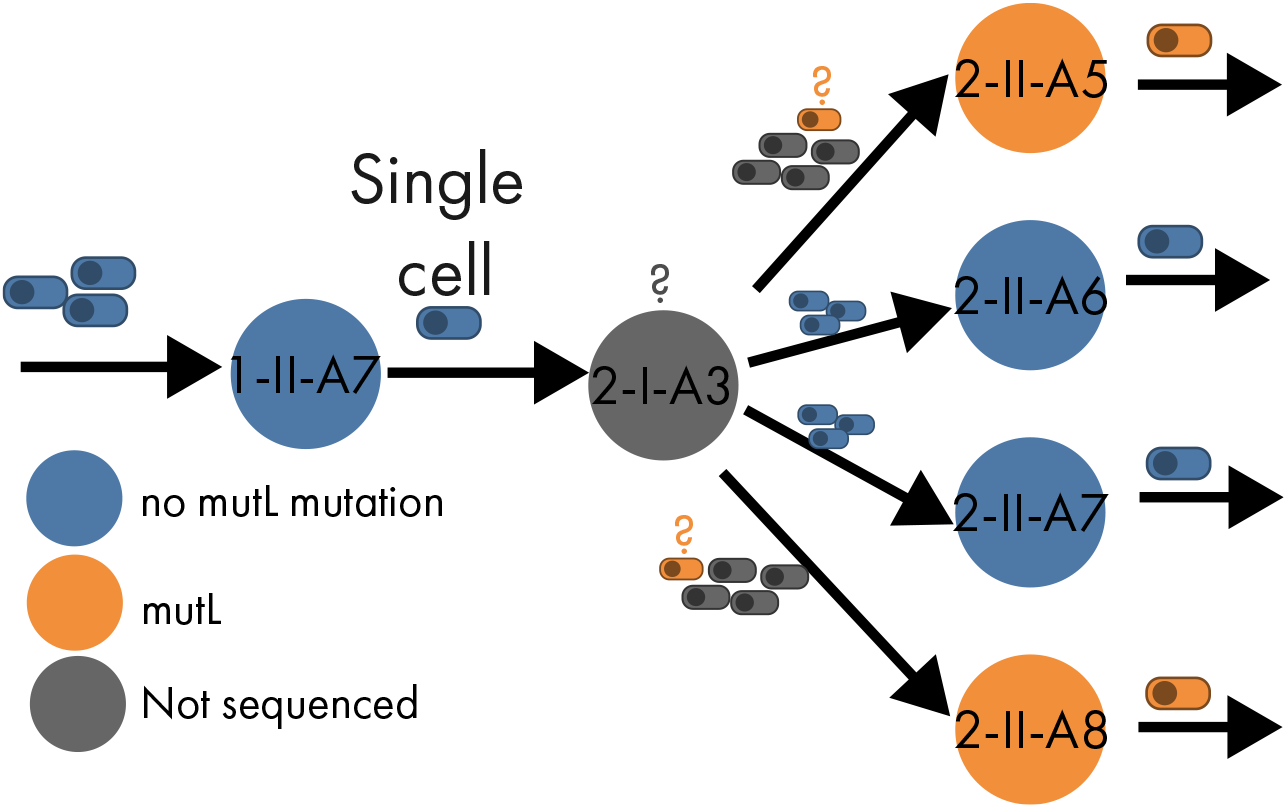
Sequencing result around 2-II-A5.

**Figure S20.**
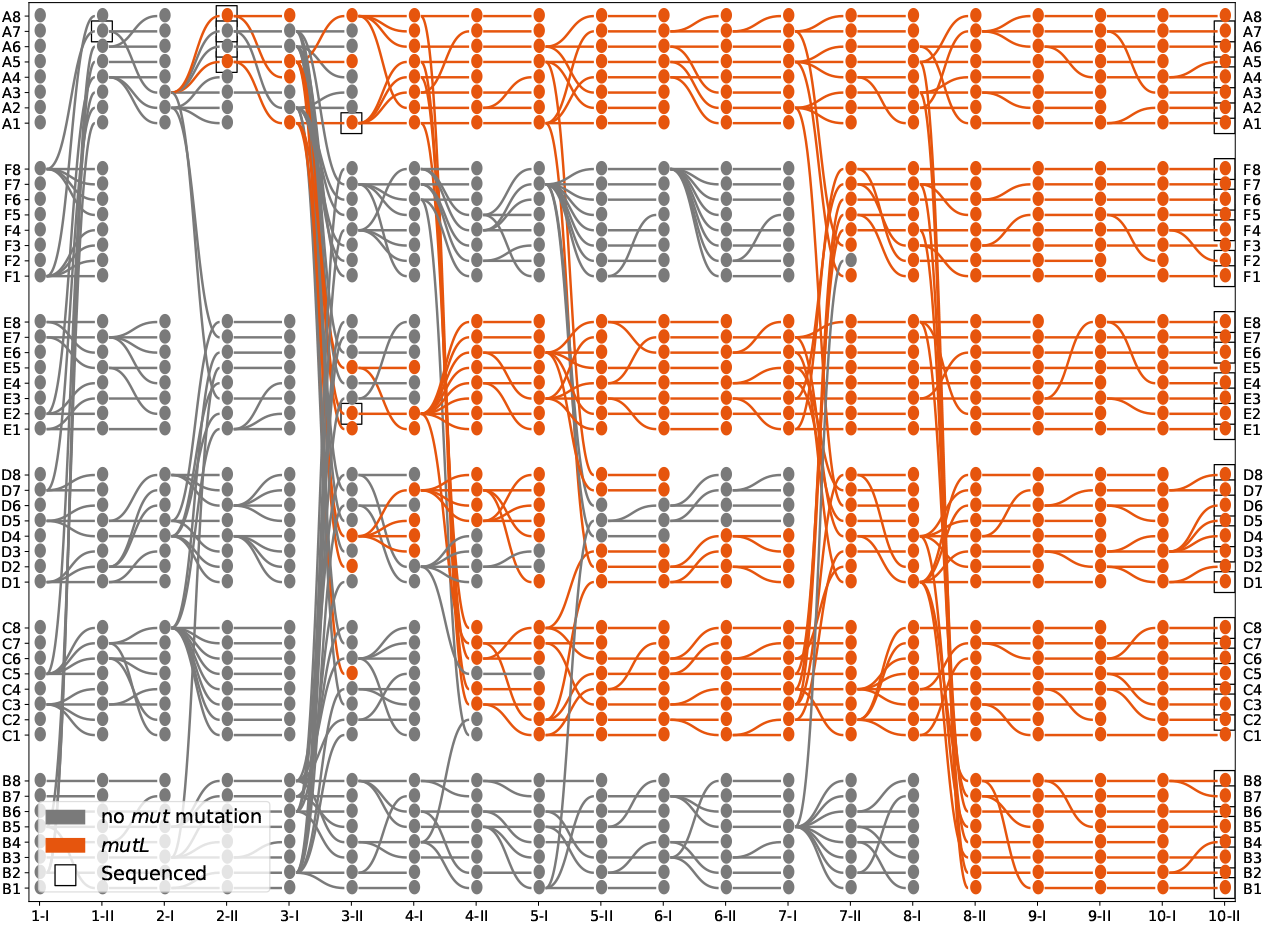
Origin of the *MutL* L103R mutation, estimated using the sequencing data from both the endpoints and within the genealogy.

**Figure S21.**
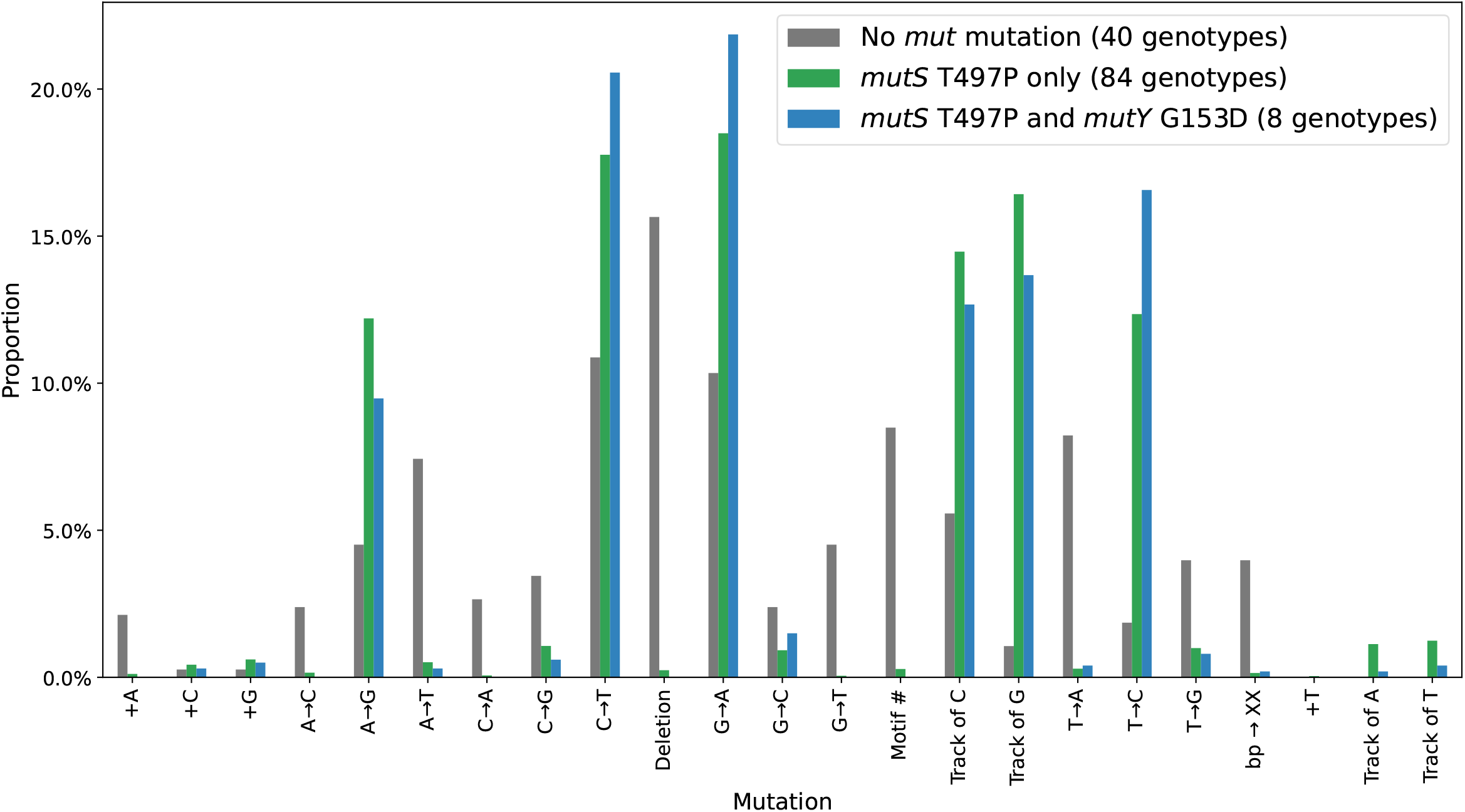
Kind of mutations in microcosms with no mut mutation, with only *mutS* T497P and with both *mutS* T497P and *mutY* G153D mutations.

**Figure S22.**
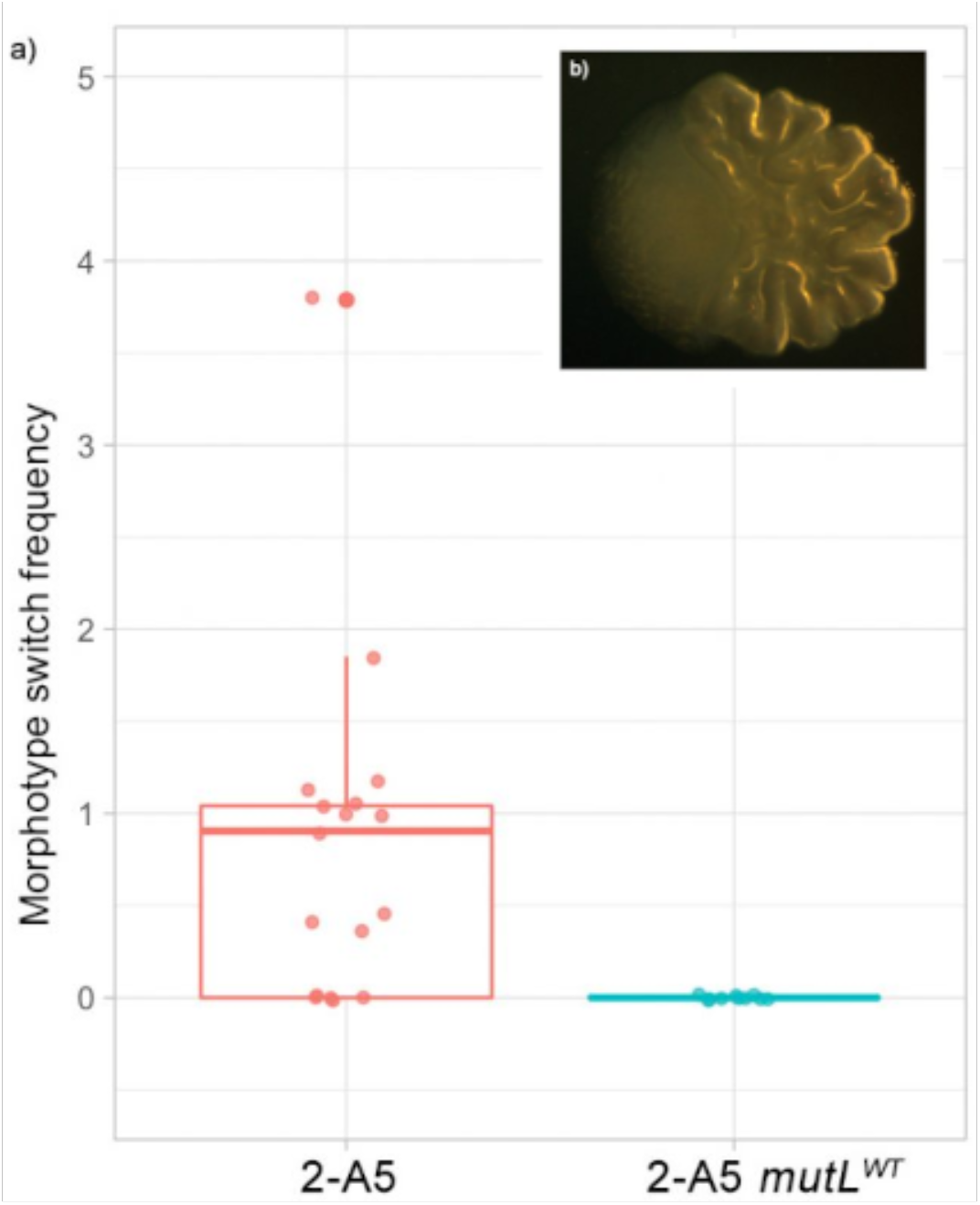
Colony morphology in 2-A5. Frequency of sectored colonies on plates (100-300 CFUs, 48h incubation) with switching *mutL* (L103R) morphotype in 2-A5 and 2-A5 with reconstituted *mutL* a); representative example of sectored colony showing morphotypic switch b).

